# The evolution of morphological castes under decoupled control

**DOI:** 10.1101/2023.02.21.529356

**Authors:** Lewis Flintham, Jeremy Field

**Author notes:** Lewis Flintham. Email: < >. Jeremy Field. Email: < >. <u>Mailing address:</u> Jeremy Field, Centre for Ecology and Conservation, University of Exeter, Penryn Campus, Penryn, Cornwall TR10 9EZ, United Kingdom.

## Abstract

Eusociality, where units that previously reproduced independently function as one entity, is of major interest in evolutionary biology. Obligate eusociality is characterised by morphologically differentiated castes and reduced conflict. We explore conditions under which morphological castes may arise in the Hymenoptera and factors constraining their evolution. Control over offspring morphology and behaviour seem likely to be decoupled. Provisioners (queens and workers) can influence offspring morphology directly through the nutrition they provide, while adult offspring control their own behaviour. Provisioners may, however, influence worker behaviour indirectly if offspring modify their behaviour in response to their morphology. If manipulation underlies helping, we should not see helping evolve before specialised worker morphology, yet empirical observations suggest that behavioural castes precede morphological castes. We use evolutionary invasion analyses to show how the evolution of a morphologically differentiated worker caste depends on the prior presence of a behavioural caste: specialist worker morphology will be mismatched with behaviour unless some offspring already choose to work. A mother’s certainty about her offspring’s behaviour is also critical – less certainty results in greater mismatch. We show how baseline worker productivity can affect the likelihood of a morphological trait being favoured by natural selection. We then show how under a decoupled control scenario, morphologically differentiated castes should be less and less likely to be lost as they become more specialised. We also suggest that for eusociality to be evolutionarily irreversible, workers must be unable to functionally replace reproductives and reproductives must be unable to reproduce without help from workers.

## 1. Introduction

Eusociality, where some individuals in a social group forfeit their own reproduction to help others to reproduce, is a phenomenon of major interest in evolutionary biology. Division of labour is most extreme in so-called obligate eusocial groups, where workers are not able to functionally replace reproductives and reproductives cannot produce reproductive offspring without help from a worker caste (see also Crespi and Yanega, 1995). These groups are characterised by morphologically differentiated castes and reduced conflict over reproduction (Bourke, 2011; Quinones and Pen, 2017; Boomsma and Gawne, 2018). In contrast, in facultatively eusocial groups, either reproductives cannot produce reproductive offspring without help from a worker caste (Crespi and Yanega, 1995), or workers are not able to functionally replace reproductive roles, but not both. In this instance, caste differences tend to be mainly behavioural (Boomsma and Gawne, 2018), and the functional constraints on caste members are typically the result of ecological conditions and life history traits (e.g. mating opportunities or swarm founding) rather than morphology. Finally, cooperatively breeding groups are those where workers can functionally replace reproductives and reproductives are capable of producing reproductive offspring without help from a worker caste (Crespi and Yanega, 1995). Different authors define these categories in slightly different ways, and the above working definitions are provided for clarity.

The significance of specialised castes as a means of social organisation in natural systems has long been understood (Oster and Wilson, 1978), and much previous work has focused on their implications and how they might evolve (Oster and Wilson, 1978; Bourke, 2011; Quinones and Pen, 2017; Cooper and West, 2018; Boomsma and Gawne, 2018). Morphological castes are startlingly diverse, ranging from simple differences in body size (Trible and Kronaeur, 2017; Kerr et al., 2019) to elaborate task-specific morphologies in taxa such as turtle ants, in which the head armour of soldiers is shaped so as to seal the nest entrance (Powell, 2016; Figure 1). Greater morphological specialisation tends to be found in larger groups (Bourke, 1999; Bourke, 2011; Ferguson-Gow et al., 2014). This is thought to reflect both reduced reproductive conflict, favouring the evolution of caste specific morphology when individuals control the expression of their own traits, and increased efficiency returns from task specialisation, favouring the evolution of morphological subcastes (Bourke, 1999; Bourke, 2011). But it may not always be reasonable to assume that individuals control the expression of their own traits. Here, we use a pairwise evolutionary invasion analysis (Avila and Mullon, 2023) to investigate how a morphological worker caste will evolve when control over worker phenotype is split between offspring and provisioners (individuals providing resources to offspring), such that manipulation can potentially occur.

**Figure 1.**
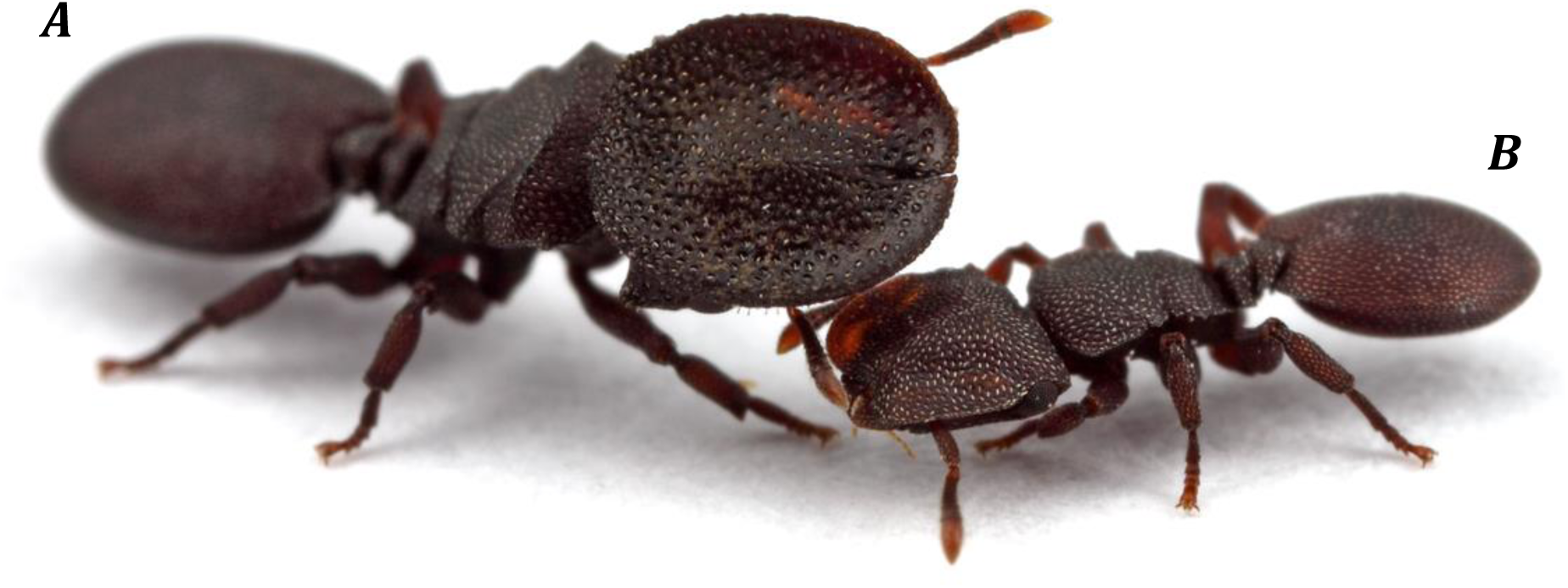
The two morphological worker castes of the turtle ant Cephalotes varians; (A) the soldier, or major worker, sporting the impressive head armour it uses to seal the nest entrance and (B) the much smaller minor worker which conducts all the foraging duties for the nest. Photo ©Alex Wild, used with permission. This image is not covered by the terms of the Creative Commons license of this publication. For permission to reuse, please contact the rights holder.

### Decoupling of control over offspring morphology and behaviour

Eusocial groups exhibiting morphological caste are characterised by a shift away from individual control. This is to say that individuals often have limited control over their own morphologies. Morphology is instead primarily determined by either the queen (and if present, the King) or the workforce as a whole (Wade, 2001; Schwander et al., 2008; Korb, 2008; Russell and Lummaa, 2009). This is most clearly seen in the social insects, in particular the bees, wasps and ants, which include a significant majority of eusocial organisms. These taxa have a holometabolic life history, in which exoskeletal development occurs in a controlled environment prior to adult emergence from the pupa (Anderson, 1972). The controlled environment is often a nest where young are provisioned and cared for by the mother or workforce (Peters et al., 2017). The use of a nest is ancestral for all social Hymenoptera (Boomsma and Gawne, 2018).

Combined with a holometabolic life history, nests give provisoners a huge amount of control over the development of immature offspring by enabling them to manipulate the environment in which offspring develop and the provisions that offspring receive (Reinhold, 2002; Karsai and Hunt, 2002; Schwander et al., 2008; Kovacs et al., 2010; Brand and Chapusiat, 2012). While the importance of nest use in driving eusociality is widely accepted (Hunt, 2007; Nowak et al., 2010; Boomsma and Gawne, 2018), its role in enabling elevated maternal control over development, and hence a greater degree of maternal manipulation, has received less attention.

If we consider individuals to be fitness maximisers (Lehmann and Rousset, 2020), we would expect those controlling phenotype to manipulate it, and hence caste, to their own benefit (Michener and Brothers, 1974; Alexander 1974; Dawkins, 1976; Kapheim et al., 2015). Previous work addressing the evolution of such maternally influenced traits has focused predominantly on the ability of individuals to resist manipulation by provisioners (Keller and Nonacs, 1993; Wenseleers et al., 2003; Uller and Pen, 2011), and on how provisioners that can manipulate both offspring behaviour and morphology might influence the origin of eusociality (Craig, 1979; Gonzalez-Forero, 2015; Gonzalez-Forero and Pena, 2021; Rodrigues and Gardner, 2021; Liu et al., 2022). Our focus in this paper, however, is not the origin of eusociality or helping behaviour, but the evolution of morphological castes *per se*, and we consequently aim to characterise when manipulation of offspring phenotype by provisioners will be favoured by selection. It has previously been noted that there may be costs as well as benefits to manipulating offspring, such as manipulator error resulting in offspring death (Wenseleers et al., 2003). We aim to identify key factors which influence the costs and benefits of manipulation and provide insight into how these factors might shape the evolution of morphological castes in the social Hymenoptera. To do so we assume a state of split control, with provisioners controlling the morphology of offspring but offspring retaining control of their behavioural phenotype. We believe this assumption to be generally representative of worker traits found in the facultatively eusocial, cooperatively breeding, and solitary Hymenoptera (Hunt, 2007; Kapheim et al., 2011; Brand and Chapusiat, 2012; Kapheim et al., 2015), from which obligate eusociality evolved. The terminology surrounding eusociality can be confusing, with different authors advocating different usages. For the purposes of this paper, “behavioural worker caste” refers to all individuals in a group that behave as workers irrespective of their morphology. The term “morphological worker caste” refers to all the individuals in a group that possess specialist worker morphology, irrespective of whether they behave as workers or reproductives. Specialist worker morphology enhances individuals’ ability to carry out worker functions while decreasing their fitness as reproductives. We consider a “worker caste” to be present in a group if there is either a behavioural worker caste or a morphological worker caste.

We will start by considering a singly mated, solitary, female hymenopteran. We assume that this female controls the morphology of her offspring, but that her offspring control their own behaviour, so that control over behaviour and morphology is decoupled. To begin with, we will also assume that any individuals which become workers will remain workers (behavioural caste is immutable) and that the mother produces an equal sex ratio. In this situation, a simple implementation of Hamilton’s rule (Hamilton, 1964; Grafen, 1985) indicates that a focal daughter, being equally related to her siblings and her offspring (r=0.5), would favour becoming a worker when she would be more productive as a worker than if she dispersed as a reproductive. In contrast, because the relatedness of a mother to her offspring (r=0.5) is twice her relatedness to her grand-offspring (r=0.25), we would expect the mother to favour her daughter becoming a worker, and hence her imposing a worker caste through altering the environment she provides for offspring development, when workers are only half as productive as reproductives. The productivity (number of offspring produced over a lifetime) of a worker relative to a reproductive is therefore critical for the arisal of a worker caste (Charnov, 1978; Bourke and Ratnieks, 1999; Reuter and Keller, 2001) and we would expect the mother to favour imposing a worker caste – perhaps by manipulating daughter morphology – before her daughter would favour being a member of it, i.e. at a lower worker/reproductive productivity ratio (Michener and Brothers, 1974; Alexander 1974; Dawkins, 1976; Craig, 1979; Kapheim et al., 2015). Inducing morphological specialization in daughters, so that they themselves will be more productive if they choose to become workers, is an obvious mechanism by which mothers could impose a worker caste. Accordingly, under our assumed state of decoupled control, we might expect the first workers to be morphologically specialised. While some authors argue that this is sometimes indeed the case (Piekarski et al., 2018; Boomsma 2022 Box 6.2), others consider that behavioural castes preceded the evolution of morphological castes in the social insects (Bourke, 2011; Quinones and Pen, 2017; Beekman and Oldroyd, 2019; Bourke 2023)), suggesting that the evolution of a morphological worker caste is somehow reliant on the prior presence of a behavioural worker caste (see Discussion and Supplementary materials Part 1).

### Productivity in the absence of specialist morphology

Across the social Hymenoptera, worker productivity varies widely (Michener, 1964; Jeanne et al., 2022), depending on factors such as group size (Michener, 1964; Jeanne et al., 2022), time constraints (Lucas and Field, 2013) and environmental constraints (Bourke, 1999; Jeanne et al., 2022) due to seasonal variations in resource availability (Poitrineau et al., 2009). Variation in worker productivity can occur between species (Field and Toyoizumi, 2020) but also within them over time (Michener, 1964; Jeanne et al., 2022). The increase in worker productivity that a specialist worker morphological trait represents for a given worker will often depend on baseline worker productivity (worker productivity without that morphological trait). To illustrate, consider a small worker whose primary function is nest defence. If a large worker is twice as likely to successfully repel an attack on the nest, then as the frequency of attacks increases, the difference in number of attacks successfully repelled by a large and small worker will also increase. We might then expect large worker size to be most likely to evolve when attacks are most frequent, i.e. when workers engaging in nest defence would be most productive.

### Summary of aims

To summarise, there is significant evidence in the nest building Hymenoptera that provisioners can control the morphology of offspring (Reinhold, 2002; Karsai and Hunt, 2002; Schwander et al., 2008; Kovacs et al., 2010; Brand and Chapusiat, 2012). Under the additional assumption that individuals control their own behaviour, we aim to characterise some of the costs and benefits determining when provisioners would favour manipulating the morphology of offspring. Drawing on the relationship between form and function, we propose a simple model illustrating the relationship between the evolution of morphological caste and the certainty the individual controlling morphology has over the behaviour of individuals occupying that caste. We then go on to use this model to explore how worker productivity in the absence of specialist morphology might influence the benefit that morphology provides, and hence the likelihood of that morphology evolving. Finally we extend our perspective of decoupled control to the evolution of eusociality more generally, with particular reference to the topics of reversibility and the arisal of helping behaviour via maternal manipulation.

## 2. The Model

We imagine a bivoltine life cycle such as that of many temperate zone cooperatively breeding bees and wasps (e.g. Seger, 1983; Field et al., 2010; Couchoux & Field 2019). A single female reproductive founds a nest then provisions a first brood of offspring (B1) herself. A fraction of the female offspring from this B1 brood will immediately disperse as reproductives and found new nests, while the remainder stay and help raise a second brood (B2) of their mother’s offspring. The females from the second brood all disperse as reproductives. Daughters from B1 which stay and help represent a behavioural worker caste. While in many small colony cooperatively breeding bees and wasps, behavioural castes are mutable, with worker takeover being common, we will start by assuming that they are immutable and that once an individual becomes a worker it will remain a worker. We formulate the following models in the style of a pairwise evolutionary invasion analysis where we ignore non-selective effects and we make the additional simplifying assumption that mutants are rare (Avila and Mullon, 2023). In addition, since we are primarily concerned with when a given manipulation trait can first be expected to spread, we focus on finding invasion conditions as opposed to evolutionary stability. A more formal derivation of the equations in this section can be found in Supplementary materials Part 4

### 2.1. The role of dispersal

We wish to determine when it will be beneficial for a mother (queen) to impart a novel morphological trait 𝒎 onto a daughter in B1, for example through the nutrition she provides the daughter during immature development. We are primarily interested in traits that provide a fitness benefit to the mother when imparted on offspring of one behavioural caste but a fitness cost to the mother when imparted to the other caste. Examples might be enlarged ovaries – benefitting reproductives but representing wasted investment on workers - and sterilisation – detrimental to reproductives but beneficial to the mother when imparted to workers as it avoids colony provisions being used to raise the offspring of workers (with which the mother shares lower relatedness). We model the expected fitness (copies of genetic material contributed to a future generation) per unit investment a mother will receive from a given daughter in the presence 𝒇^′^ and absence 𝒇 of morphological trait 𝒎 (Table 1). It may be helpful to envisage trait 𝒎 as a change in offspring morphology resulting from a mutant allele for manipulation in the mother. 𝒇^′^ and 𝒇 then represent the fitness a mother, with (𝒇^′^) and without (𝒇) this mutant allele, will expect to receive from a given daughter.

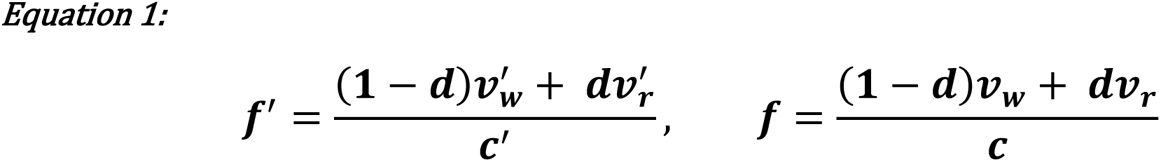

**Table 1.** Parameters used in our models.

| | With morphological trait $m$ | Without morphological trait $m$ (“baseline value”) |
| --- | --- | --- |
| Fitness a worker daughter will provide to the mother | $v'_w$ | $v_w$ |
| Fitness a reproductive daughter will provide to the mother | $v'_r$ | $v_r$ |
| Cost to mother of producing a daughter | $c'$ | $c$ |
| Probability a daughter disperses (independent of morphology) | $d$ | $d$ |
| Expected fitness per unit investment a daughter will provide to the mother | $f'$ | $f$ |
| Probability a daughter disperses when trait $m$ increases the likelihood of daughters staying as workers. | $d(1 - a)$ | $d$ |

Trait 𝒎 will be expected to invade when;

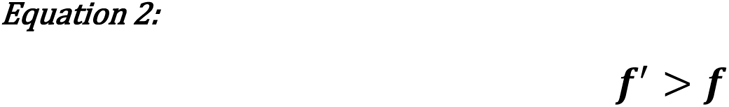

Thus the expected fitness per unit investment a mother will receive from a given daughter is dependent on the probability of that daughter dispersing as a reproductive 𝒅, as well as on the cost of producing the daughter. Since 𝒎 here is considered to be a novel trait, we assume that daughters do not respond to the presence of the trait behaviourally, and have the same probability of dispersing with or without the trait (Table 1). We will later relax this assumption to see how morphology will evolve when offspring do change their behaviour in response.

Substituting Equation 1 into Equation 2 we obtain the parameter space where we expect imparting trait 𝒎 to be selected for (Figure 2):

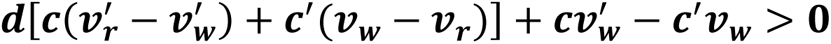

**Figure 2.**
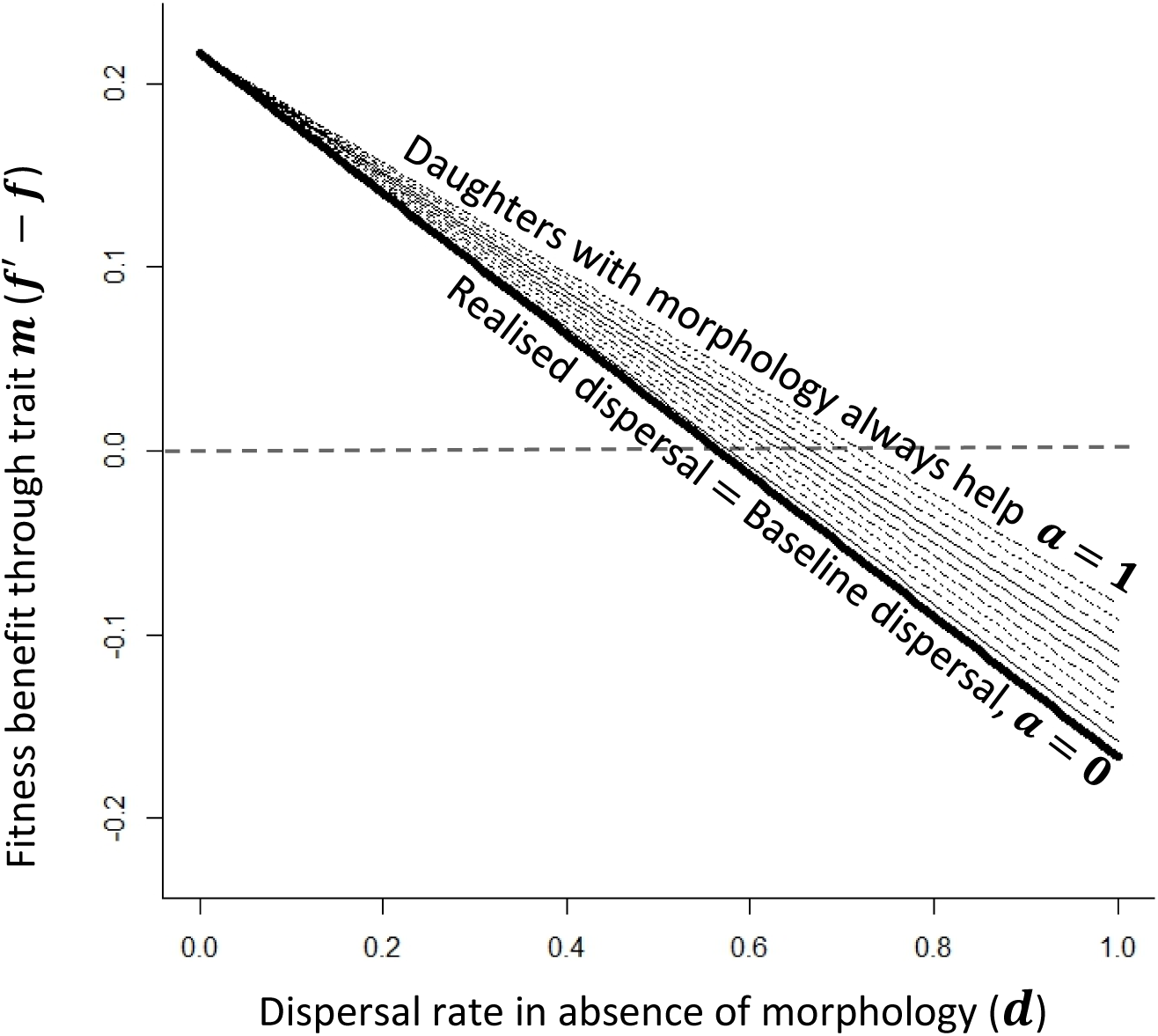
The linear relationship between the net fitness increase a provisioner will receive from imparting morphological trait 𝒎 onto a daughter (𝒇^′^ − 𝒇), and the baseline probability of a daughter dispersing (see Table 1 for parameter definitions). We have plotted this for 𝟎 ≤ 𝒂 ≤ 𝟏 at 0.1 increments. The thick line represents 𝒂 = 𝟎 and hence Equation 3. For parameter values 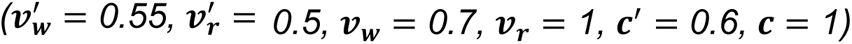. Dashed line at y=0 represents the threshold for manipulators imparting trait 𝒎 to invade (Equation 2)

When 𝒎 is a trait conferring specialist worker morphology, we would expect it to increase the fitness the mother receives from workers per unit cost more than the fitness she receives from reproductives per unit cost 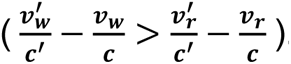 . When no daughters from B1 disperse as reproductives (𝒅 = 𝟎), a trait which increases the fitness a mother receives from a worker per unit cost 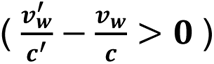 will always be selected for. More generally, there will be a negative linear relationship between probability of dispersal and the likelihood of manipulation for trait 𝒎 being favoured (Figure 2): the smaller the proportion of B1 daughters that disperse as reproductives, the more likely that it is advantageous for a mother to impart worker morphology, because morphology will less often be mismatched with behaviour. Because we are primarily interested in worker morphology which decreases the fitness per unit cost of a reproductive daughter, and behaviour here is considered independent of morphology, we do not expect such specialist worker morphology to evolve without the prior evolution of helping behaviour: if all B1 daughters become reproductives, trait 𝐦 inevitably reduces offspring fitness.

### 2.2. Behaviour as a response to morphology

In the previous section, morphological evolution responds to behaviour in the sense that worker morphology cannot evolve without the prior presence of a behavioural worker caste. Next, we consider the possibility that individuals can respond plastically to their morphology (El Mouden and Gardner, 2008) by choosing to disperse as reproductives or not. Morphological traits may not always be novel, and if we assume that trait 𝒎 was historically present in our population, it seems likely that B1 daughters will have evolved to alter their dispersal decisions in response to the trait’s presence. To model this new situation, we introduce parameter 𝒂 (Table 1; Figure 3), which represents the effect of trait 𝒎 on the probability of B1 daughters dispersing.

**Figure 3.**
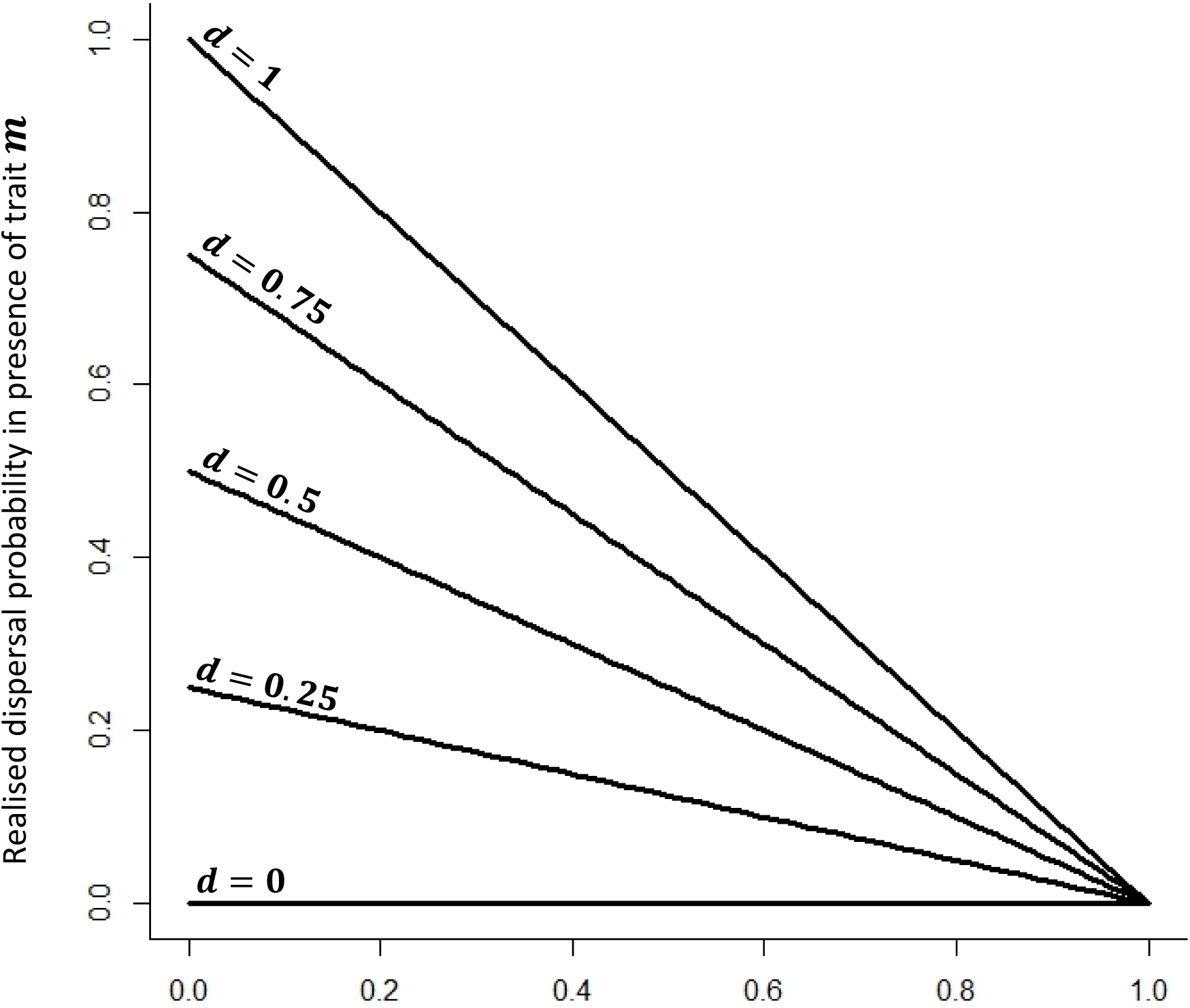
The relationship between 𝒂, the degree to which daughters change their dispersal behaviour in the presence of trait 𝒎, and realised offspring dispersal. Different lines represent different values of baseline dispersal (dispersal in the absence of trait 𝒎; 𝟎 ≤ 𝒅 ≤ 𝟏) in 0.25 increments. When 𝒂 > 𝟎 daughters are more likely to work (Table 1).

When 𝒂 > 𝟎, daughters with trait 𝒎 are more likely to stay as workers, and when 𝒂 = 𝟏 they always stay. Conversely, when 𝒂 < 𝟎, daughters with trait 𝒎 are more likely to disperse as reproductives - this area of the parameter space has limited relevance to the evolution of specialist worker traits, and is explored in Supplementary materials Parts 2 and 3.

Incorporating parameter 𝒂 gives us the following equation:

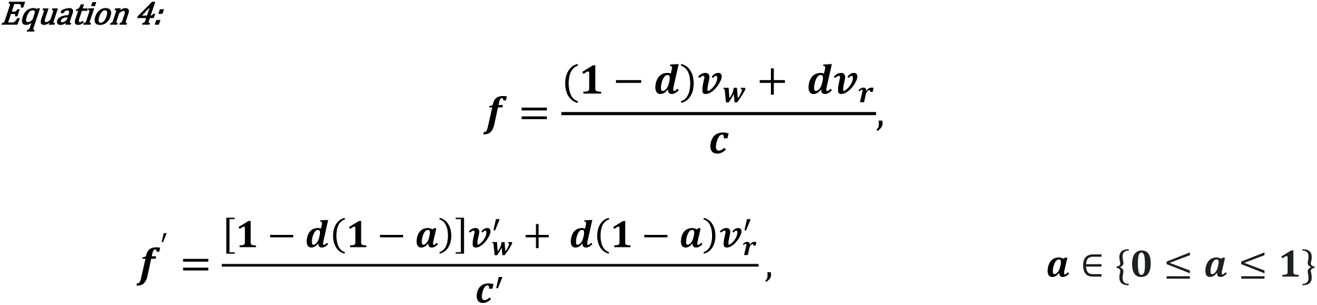

When 𝒂 = 𝟎, the presence of trait 𝒎 has no effect on the dispersal decisions of daughters, and Equation 4 reduces to Equation 1. When daughters can respond to the presence of morphological traits (𝒂 ≠ 𝟎), mothers obtain a degree of indirect control over daughter behaviour through manipulating their morphology. The extent of this control is represented by the magnitude of 𝒂. In order to determine when imparting 𝒎 will be favoured by selection now that daughters can respond to it behaviourally, we can substitute Equation 4 into Equation 2 and rearrange to give the parameter space where we expect manipulation for trait 𝒎 to invade as:

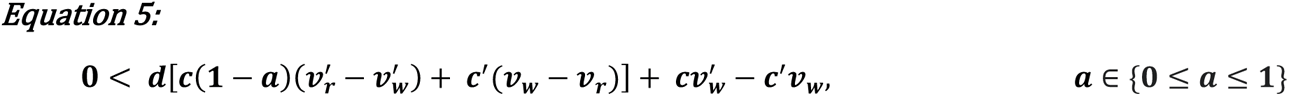

We again find a linear relationship between 𝒅, the baseline probability of dispersal, and the likelihood of manipulation for trait 𝒎 invading (Figure 2). The relationship is again negative when 𝒎 confers specialist worker morphology, but the gradient is now dependant on the value of 𝒂, such that 𝒂 is moderating the relationship between baseline dispersal and the likelihood of manipulation for trait 𝒎 invading (Figure 2). This is because 𝒂 causes the realised dispersal choices of daughters to correlate with their morphologies, altering the degree of mismatch between form and function. Parameter 𝒂 therefore introduces a mechanism through which mothers can use their control over daughter morphology to influence daughter behaviour, and consequently moderate the certainty a mother has about how an individual daughter will behave.

To take an example, trait 𝒎 might represent a simple reduction in the size of B1 daughters. Smaller daughters will be cheaper to produce (Brand and Chapusiat, 2012), so that if worker performance is independent of size (e.g. Couchoux and Field, 2019), the fitness per unit investment received by a mother from a worker will increase. However, we might also expect trait 𝒎 to decrease the fitness per unit investment from daughters that disperse as reproductives: reproductives must found new nests and lay eggs, and smaller reproductives may perform less well with these tasks (Couchoux and Field, 2019). If there has historically been variation in the size of daughters independent of trait 𝒎, perhaps environmentally induced, it seems plausible that daughters will have already evolved to alter their dispersal decisions according to their size, with smaller daughters being more likely to stay as workers (Brand and Chapusiat, 2012). Our value of 𝒂 in this instance would be positive (since daughters with trait 𝒎 are more likely to stay as workers; Supplementary materials Part 3), and manipulation for trait 𝒎 will be favoured by selection over a larger range of values of 𝒅 than if daughters are unable to respond plastically to morphology (see Figure 2).

We can also extend the idea of maternal certainty over offspring behaviour to instances where workers have the possibility of inheriting the queen role. This changes the proportion of time a mother expects a daughter to spend as a worker or reproductive for a given probability of dispersal (Toyoizumi and Field, 2014). Time spent as an inheriting worker is assumed equivalent to time spent as a reproductive in the sense that the optimal morphology in both cases is reproductive morphology (Supplementary materials Parts 1 and 2). Thus the likelihood of a mismatch between form and function when a mother imparts specialist worker morphology onto a daughter increases, making specialist worker morphology less likely to be favoured. However, since time spent as a reproductive or inheriting worker still depends on the probability of dispersal (𝒅), the qualitative nature of the relationship between dispersal and the parameter space where we might expect to see the evolution of specialist morphology remains similar (Figure S1).

We note that when 𝒂 > 𝟎, the sex ratio of dispersing offspring that a mother with the trait for manipulation produces will be more male biased than the sex ratio of the offspring produced by a mother without the trait: some of the female offspring which would have dispersed now stay and help. This will introduce an element of frequency dependence that is not taken into account in our analysis and would manifest as changes in 𝒗_𝒓_ relative to 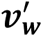. The more common the manipulation trait becomes, the more male biased the mating pool, lowering the reproductive value of male offspring and hence the expected fitness of a mother who manipulates her offspring relative to one who does not. However, so long as the population has a sufficiently large, well mixed mating pool and an evolutionarily stable sex ratio, this will not impact the invasion conditions for a rare manipulation trait (Avila and Mullon, 2023).

### 2.3. Trait benefit as a function of baseline productivity

The fitness represented by a worker offspring bearing trait 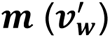 is likely to be correlated with the fitness represented by a worker that lacks trait 𝒎 (𝒗_𝒘_): trait 𝒎 represents a single modification of worker morphology, leaving the remaining morphology unchanged. Thus, when the productivity of a worker without trait 𝒎 changes we expect the productivity of workers with trait 𝒎 to change as well, so that selection on trait 𝒎 will depend on 𝒗_𝒘_. Now it may be the case that the fitness benefit conferred by trait 𝒎 varies with 𝒗_𝒘_ such as in the relationship 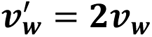, where trait 𝒎 doubles the fitness represented by a worker. Again taking the example of worker size, this would be the case if a large worker is twice as productive as a small worker. The difference in productivity between large and small workers (the fitness benefit of being large) would then be equal to the productivity of a small worker. We use changes in productivity and fitness interchangeably here, but note that the change in a worker’s productivity caused by a worker trait may not always be equivalent to the change in the mother’s fitness, if the offspring produced are not all equivalent (e.g. if larger workers invest less into producing their own male offspring). Even when changes are equivalent, this may not be apparent in the field, for example if immature or adult workers with a morphological trait have higher mortality, but the productivity of only surviving workers is measured (Cant and Field, 2001; Couvillon and Dornhaus, 2010). Substituting 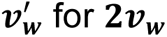 for 𝟐𝒗_w_ in Equation 5 we can rearrange to give the following:

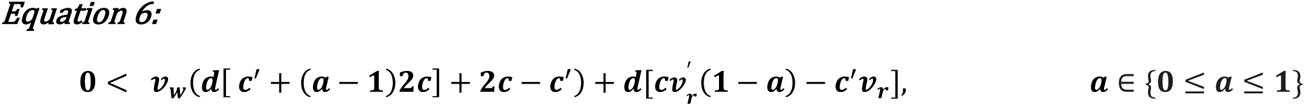

As discussed above, when 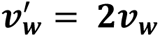 we tend to see a strong positive relationship between 𝒗_𝒘_ and the likelihood that manipulation for trait 𝒎 will invade (Figure 4). However, Equation 6 also highlights how this is not always the case. For example, since it is the benefit per unit cost which primarily determines whether a trait will be selected for, if the cost of producing a daughter with trait 𝒎 (𝒄^′^ − 𝒄) is very large (𝒄^′^ > 𝟐𝒄), then we can still generate negative relationships between 𝒗_𝒘_ and the likelihood that manipulation for trait 𝒎 will invade, despite setting 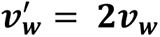 (Supplementary materials Part 3).

**Figure 4.**
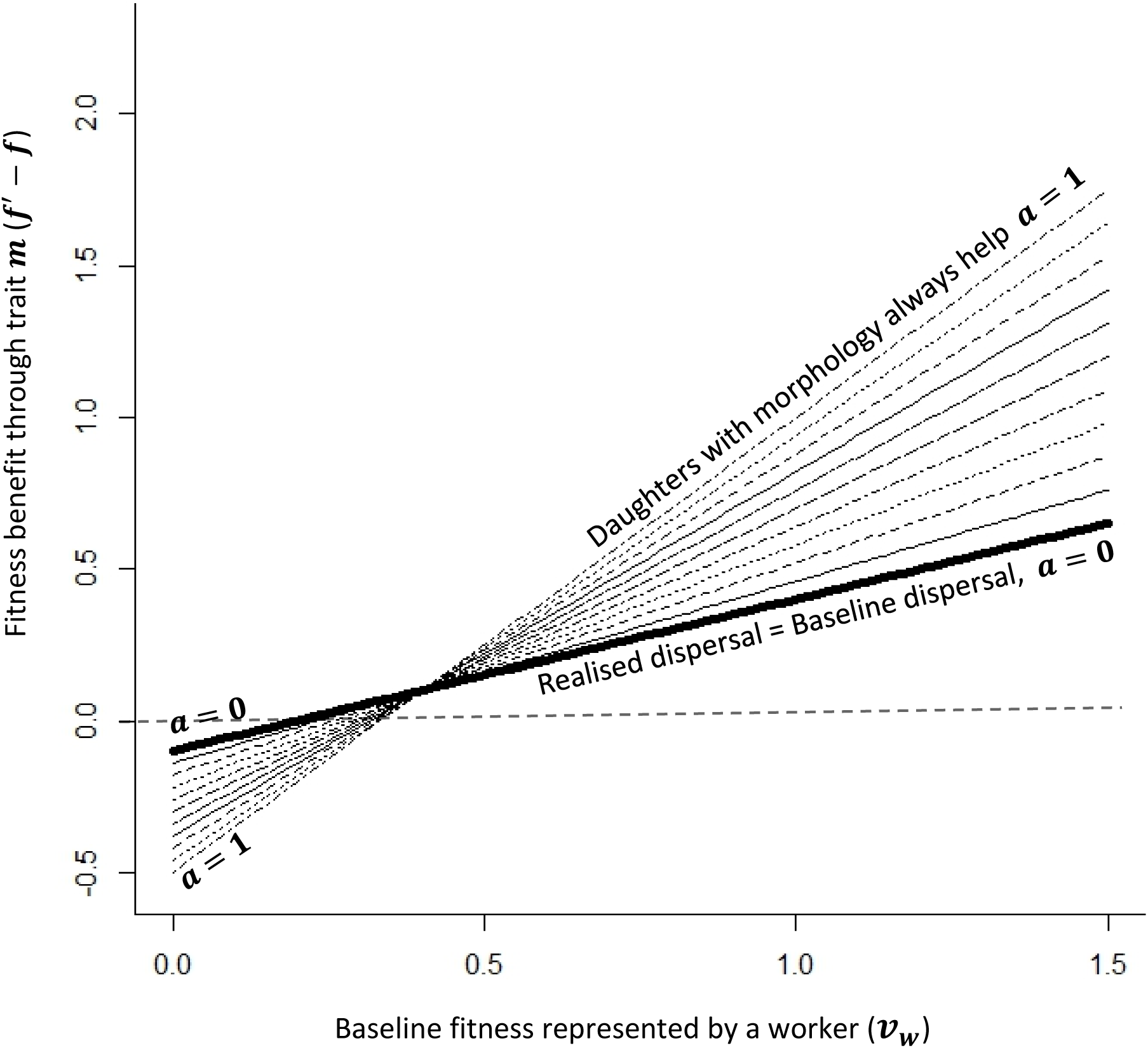
The relationship between the fitness of a worker in the absence of a trait and the likelihood of that trait being selected, when workers with that trait are twice as productive as those without it 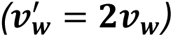. We have plotted this for 𝟎 ≤ 𝒂 ≤ 𝟏 at 0.1 increments with parameter values 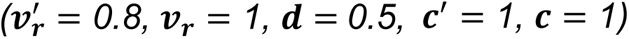. Dashed line at y=0 represents the threshold for when manipulators imparting trait 𝒎 will invade (Equation 2).

More generally we find that if there is a positive relationship between the productivity benefit of a specialist worker trait and baseline worker productivity; so long as the increased benefit provided by the trait remains proportionally greater than the cost of producing it, and if the trait does not significantly increase the likelihood of daughters dispersing, an increase in baseline worker productivity will lead to increased selection for that trait (Supplementary materials Part 3). This effect is not dependent on decoupled control (Supplementary materials Part 2).

## 3. Discussion

Our analysis shows that unless a behavioural worker caste has already evolved, a new morphological trait that increases worker fitness but decreases the fitness of reproductives cannot invade, for the simple reason that all offspring will be reproductives. This may seem obvious, but it means that although we might expect queens to use their control over offspring morphology to impose worker behaviour at a lower threshold benefit/cost ratio than the threshold at which offspring themselves would work voluntarily, worker behaviour will not evolve unless the worker’s own threshold has already been reached. Thus, contrary to some previous suggestions (Michener and Brothers, 1974; Alexander 1974; Craig, 1979; Kapheim et al., 2015; Nonacs and Denton 2023), maternal manipulation of offspring morphology cannot induce helping behaviour where it was not previously present: the origin of helping is instead constrained by the productivity threshold at which offspring themselves will choose to work voluntarily. We note that in Kapheim et al.’s (2015) model, daughters are assumed to respond to morphological manipulation by staying at their natal nests and helping more frequently i.e. plasticity for helping behaviour is assumed to already be present in the population, and must have evolved historically (as in our second model when 𝒂 > 0).

Our analysis thus suggests that behavioural castes should always precede morphological castes in the social Hymenoptera, and that the evolution of morphological castes relies on provisioners having a high degree of certainty over which offspring will help. If morphological manipulation (of worker traits) cannot be limited to those offspring which are less likely to disperse as reproductives, morphological castes would not be expected to evolve (Supplementary materials Part 4). We do not attempt a systematic review here, but several empirical patterns seem broadly consistent with this. First, behavioural castes are often observed when no morphological caste differentiation has been reported, whereas the opposite is not true. While the possibility that there are subtle morphological differences between castes has not been examined in most apparently facultatively eusocial and cooperatively breeding taxa, there are clear examples of behavioural castes exhibiting no size or other morphological differentiation (e.g. Field 2008; Field et al., 2010). Second, we never see extreme morphological caste specialisation when offspring have a low likelihood of helping, but we do sometimes see limited morphological caste specialisation when offspring have a high likelihood of helping (O’Donnell and Jeanne, 1993; Tibbetts, 2006; Toth et al., 2009), consistent with the evolution of morphological castes being reliant on provisioners having a high degree of certainty of offspring behaviour. An additional pattern we might expect to see is that taxa with morphologically differentiated castes should be nested within lineages that exhibit only behavioural castes. Here, the evidence is mixed. Allodapine bees may represent an example of the predicted pattern (Schwarz et al., 2007), but it has been argued that the phylogeny of vespid wasps indicates that eusociality and morphological castes evolved simultaneously (Piekarski et al., 2018; see also Boomsma 2022 Box 6.20). Our analysis suggests that this is unlikely. One possibility is that morphologically undifferentiated ancestral taxa are now extinct and therefore absent from phylogenetic reconstructions, as is presumably the case for ants, all of which are eusocial. Another possibility is that once unmanipulated offspring began to help, selection favoured mothers conferring morphological specializations on them, which in turn favoured manipulated offspring being even more likely to help (as in our second model, and discussed in Supplementary materials Part 3). The resulting feedback loop could make helping and morphological specialization appear to have evolved together.

We have primarily considered queens as being the provisioners controlling the morphology of offspring. However, once a behavioural caste has been established as we see in the social Hymenoptera, it becomes increasingly common for the workforce as a whole to instead control offspring morphology (Cameron and Robinson, 1990; Inoue et al., 1996; O’Donnell and Jeanne, 1993; Giray et al., 2005). The worker productivity at which the workforce favours offspring helping will depend on the number of times the queen has mated and also on the sex ratio of offspring produced (Reuter and Keller, 2001). However, we would still expect to see a disparity between workforce and offspring over the worker productivity at which helping is preferred (Stubblefield and Charnov, 1986).

More importantly, our findings suggest that in the social Hymenoptera, where it is likely that provisioners control offspring morphology while offspring control their own behaviour (Reinhold, 2002; Karsai and Hunt, 2002; Kapheim et al., 2011; Kovacs et al., 2010; Brand and Chapusiat, 2012; Kapheim et al., 2015), the certainty that provisioners have about offspring dispersal decisions is a key constraint on the evolution of increasingly differentiated morphological castes (Equation 3). Reduced certainty results in greater mismatch between form and function. These conclusions are expected to hold even when individuals can switch behavioural castes within a lifetime (Supplementary materials Part 2). It is important to emphasise that the certainty provisioners have over offspring dispersal decisions can be increased through mechanisms other than a direct offspring response to morphology, for example where there is seasonal variation in offspring dispersal. A clear example of this is in temperate cooperatively breeding bees and wasps, where daughters from the first brood (B1) are more likely to become helpers than daughters from the second brood (B2), which always disperse as reproductives (section 2.1). Provisioners therefore have greater certainty about daughter behaviour, and parental manipulation of offspring morphology might evolve if it could be limited to first brood offspring. The ability for offspring to adjust their dispersal decisions as a response to their morphology can also give provisioners a significant degree of indirect control over the behaviour of female offspring and allow morphological traits to be selected for at lower baseline rates of dispersal than we might otherwise expect (Equation 5).

### Baseline worker productivity can affect the likelihood of specialist morphology evolving

We have shown that the baseline productivity of workers (in the absence of specialised morphology) can be important in determining when specialist morphology is likely to evolve. The gain in worker productivity (and hence provisioner fitness) provided by a morphological trait may vary as a function of baseline worker productivity, an example being a trait that increases worker size, where larger workers are always twice as productive as smaller workers. In this case, when smaller workers have a productivity value of 5, larger workers would have a productivity of 10 (5 greater), but when smaller workers have a productivity value of 7, larger workers have a productivity of 14 (7 greater). A trait causing larger worker size will then be most likely to evolve when baseline worker productivity is high, since this is when larger workers provide the largest gain. Generally, if the extra fitness provided by a specialised morphological trait increases with baseline worker productivity (linearly or otherwise), the trait is most likely to be favoured when baseline productivity is high - unless that trait is especially costly (Supplementary materials Part 3). In practice, determining the relationship between morphology and productivity can be complex, with morphological traits not always enhancing all worker tasks equally, and epistasis for fitness (Bank, 2022) often making the extraction of simple empirical correlations difficult (Grimm et al., 2005; Kugonza et al., 2011; Wen et al., 2015).

### Decoupled control, reversibility, and loss of conflict between offspring and provisioners

We now extend our analysis by considering how it might relate to the evolutionary reversibility of morphological castes. It has previously been suggested that the presence of a morphologically specialised worker caste, in particular a sterile worker caste, represents a state of irreversibility wherein the worker caste can no longer be lost evolutionarily (Boomsma 2013; Boomsma and Gawne 2018). The perspective we have taken on control structure reveals a mechanism that could underlie reduced reversibility: with decoupled control, morphological castes become less likely to be lost as they become more specialised, and conflict between offspring and provisioners over offspring caste fate may disappear (see Bourke 2011: 178 for a related discussion).

After a worker caste has evolved, it might be lost when the environment changes so that worker productivity decreases. Indeed, phylogenetic work shows that the largely behavioural worker caste seen in sweat bees has frequently been lost (Wcislo and Danforth, 1997), probably when bees entered environments where the season was too short for workers to be productive (Lucas and Field, 2013). Once a morphological worker caste controlled by provisioners is present, however, we expect workers to evolve to respond to their morphology when making decisions about their behaviour (section 2.2). Figure 5 illustrates the implications of this. For simplicity, we focus on the case when offspring with specialist worker or reproductive morphology are equally costly to produce – an example might be where worker morphology simply involves mandibles shaped specifically for defensive or foraging tasks. Initially, offspring prefer to disperse at higher worker productivities than provisioners prefer – productivities at which provisioners, because they are more closely related to the queen’s offspring than to her grand-offspring, would prefer them to work (Figure 5A; see Introduction). With each additional morphological specialisation, however, the behavioural caste that an offspring chooses should be constrained more and more by its morphology: offspring will incur larger and larger costs if they mismatch their behaviour to their morphology. The effect is almost ratchet-like: each additional morphological specialism reduces the fitness of daughters that mismatch their behaviour to their morphology by dispersing, so that worker productivity must drop progressively lower for offspring with worker morphology to nevertheless prefer dispersal (Figure 5B).

**Figure 5.**
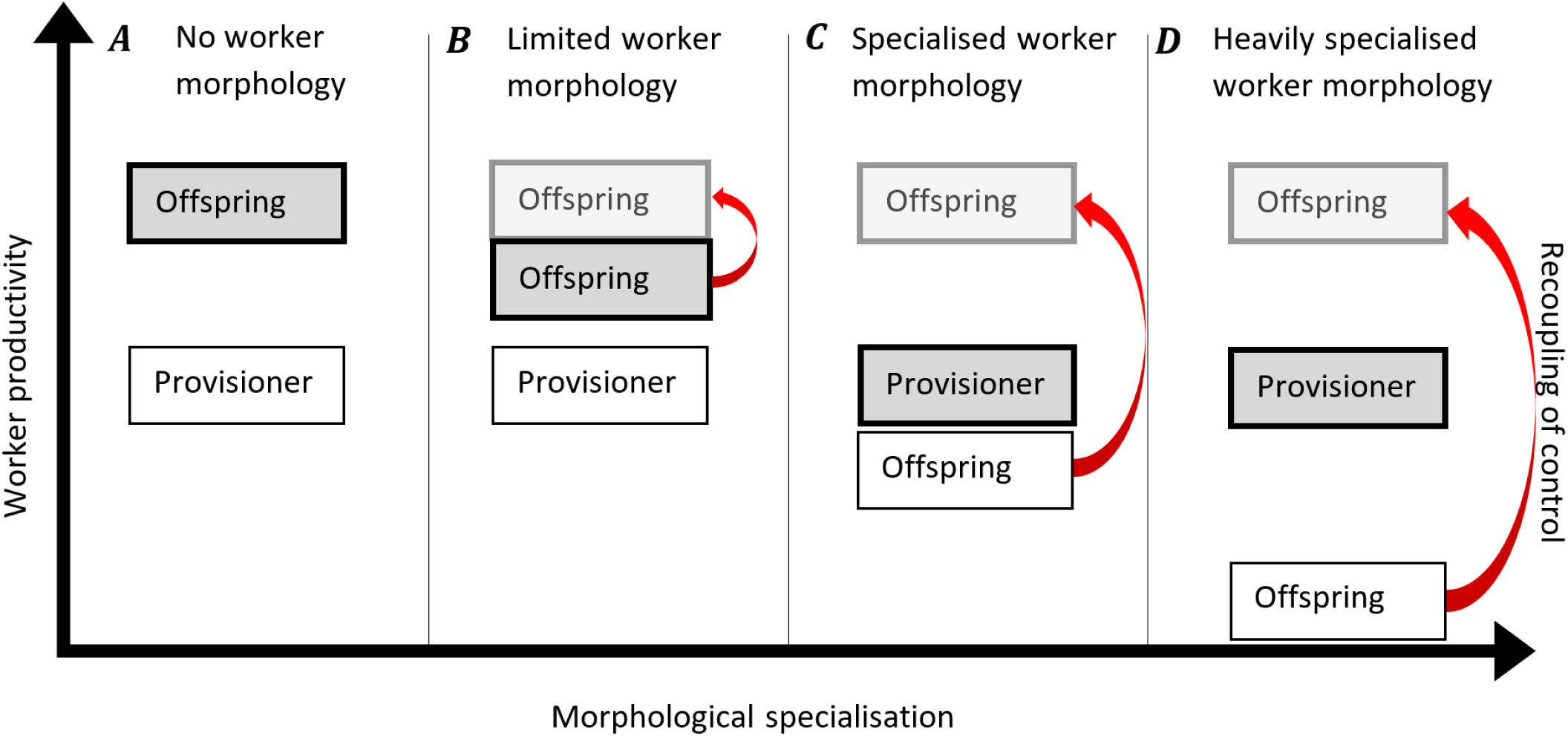
The relationship between the level of worker morphological specialisation and the threshold worker productivity at which offspring dispersal is favoured by two parties, offspring and provisioners. Preferences under decoupled control are represented by shaded (bold-edged) and unshaded boxes. The worker productivities at which offspring will begin to disperse, so that the worker caste is lost, are represented by shaded/bold-edged boxes. Red arrows linked to faded boxes indicate the possibility that offspring regain control of their morphology, recoupling it with control of behaviour, so that they do not incur mismatch costs when dispersing.

In contrast, because provisioners control offspring morphology, the worker productivity at which provisioners prefer offspring to disperse remains constant (figure 5): if the fitness that provisioners obtain from workers with specialist worker morphology falls below what they obtain from dispersers with reproductive morphology, provisioners are expected to impart reproductive morphology – offspring should respond to this by dispersing, thus avoiding any mismatch. Eventually, once morphological castes are sufficiently specialised, the ratchet-like effect will mean that the worker productivity at which offspring with worker morphology themselves prefer dispersal drops below that at which provisioners prefer their offspring to disperse (Figure 5C). It should then be provisioners, not offspring, that first cause the loss of a worker caste during evolution, should worker productivity drop sufficiently. The worker caste is then no longer ‘self-determined’ (Bourke and Ratnieks, 1999) by the offspring and can instead be considered to be ‘manipulator enforced’ by the provisioner. At this point, realised conflict between offspring and provisioners over dispersal is lost, with both parties preferring the same outcome (the dispersal of offspring).

If at any point individual offspring are able to regain control of their morphology during evolution (recoupling control over morphology and behaviour), as seems to be the case in stingless bees (Meliponini) and a few other obligately eusocial taxa (Ishay and Landau, 1972; Wenseleers et al., 2003; Kaptein et al., 2005; Mas and Kolliker, 2008), the threshold worker productivity required for a worker caste to be lost becomes wholly determined by individual offspring (returning the worker caste to being ‘self-determined’; red arrows in Figure 5). Offspring can then choose whether to work or disperse as reproductives without incurring costs through mismatching morphology and behaviour. This too represents a loss of realised conflict over dispersal, since now only one party is able to influence the outcome, this time resulting in dispersal being preferred at a higher level of worker productivity. The apparently offspring-controlled caste determination seen in the Meliponini (Wenseleers et al., 2003; Wenseleers and Ratnieks, 2004) could therefore represent a vulnerability to selection for loss of the worker caste, although so long as other aspects of meliponine life history remain, such as swarm founding that requires workers, such a reversal seems unlikely.

### Extreme worker traits and evolutionary irreversibility

It is useful to consider where extreme morphological specializations such as worker sterility might appear in the above sequence (Figure 5). The very high costs of mismatch that such specializations entail means that they should either be associated with a recoupling of control over morphology and behaviour, such that mismatch can be avoided, or with a manipulator enforced worker caste. This is because, if control has not been recoupled and mismatch cannot be avoided, we would expect less extreme specialisations, with lower costs of mismatch, to evolve before extreme traits like sterility – when provisioners are less certain about offspring behaviour. These less extreme traits might be sufficient on their own to cause the worker caste to become manipulator enforced (Figure 5C). Additional, more extreme morphological specialisations such as sterility would not then further reduce the likelihood of a worker caste being lost, since the point at which workers themselves might prefer to disperse would already have been passed (Figures 5C and 5D).

In the previous section, we suggested that the worker caste is less likely to be lost as it becomes more specialised. We here consider more generally what evolutionary ‘irreversibility’ means in the context of eusociality (Boomsma and Gawne, 2018; see also Nonacs and Denton 2023). If reproductives are dependent on help from a worker caste to produce reproductive offspring – for example if the life cycle requires swarm founding – then that worker caste (morphologically specialised or otherwise) cannot be lost evolutionarily following a reduction in worker productivity, without resulting in extinction (Crespi and Yanega, 1995). It might then seem reasonable to consider the worker caste evolutionarily irreversible at this point. Irreversibility for the worker caste is then conditional on the capabilities of individuals of the reproductive caste, not on how morphologically specialised the worker caste itself has become. However, the capabilities of individuals in the reproductive caste could change – for example if the environment changes so that swarm founding is no longer essential. Likewise, if workers are unable to functionally replace reproductives in producing both male and female reproductive offspring, so long as worker capabilities remaining unchanged the reproductive caste cannot be lost evolutionarily without resulting in extinction. Thus, both of the criteria for obligate eusociality (Crespi and Yanega, 1995) must be maintained for eusociality to be considered evolutionarily irreversible, at least at a given point in time: workers must be unable to functionally replace reproductives, and reproductives must be unable to produce reproductive offspring without help from a worker caste. We note that while being strongly correlated with morphological specialisation (Bourke, 2011; Quinones and Pen, 2017; Boomsma and Gawne, 2018), both criteria could in principle be maintained entirely through ecological conditions and life history traits, such as when workers lack mating opportunities (inability for workers to functionally replace reproductives) and swarm founding is favoured (reliance of reproductives on the worker caste to produce reproductive offspring).

### The prevalence of decoupled control

While we have focused primarily on the evolution of a morphological worker caste in the facultatively eusocial and cooperatively breeding Hymenoptera, the relationships we have discussed underlying the evolution of morphological traits will hold for any level of social organisation or study system so long as our assumptions regarding the immutability of morphological castes and split control are met (Supplementary materials Part 2). Our assumption that provisioners control the morphology of offspring may not be equally valid for all morphological traits (e.g. body size vs body shape), and must be evaluated on a trait by trait basis. Relaxing these assumptions represents a possible future extension of our ideas and would greatly generalise our results, though it seems likely that the key linear relationships we identify involving behavioural certainty and baseline productivity will remain important. Shared control seems especially likely in progressively provisioning taxa, with substantial evidence that offspring begging influences provisioning decisions in obligately eusocial Hymenoptera (Ishay and Landau, 1972; Creemers et al., 2003; Kaptein et al., 2005; den Boer and Duchateau, 2006; Mas and Kolliker, 2008; He et al., 2016). However, evidence for offspring resistance to provisioner manipulation in the Hymenoptera is limited outside of obligately eusocial species (Ishay and Landau, 1972; Wenseleers and Ratnieks, 2004; Kaptein et al., 2005; Mas and Kolliker, 2008; Gonzalez-Forero, 2015). Begging behaviours in solitary, cooperatively breeding, or facultatively eusocial progressively provisioning taxa have not to our knowledge been reported, suggesting that our assumption of queen control over morphology but offspring control of behaviour is reasonable before morphological castes evolve. It is possible, however, that the absence of reported begging outside the obligately eusocial Hymenoptera instead reflects a disparity in the volume of research conducted.

## Conclusions

1. There is evidence that in the social Hymenoptera, the evolution of a morphological worker caste is reliant on the prior presence of a behavioural worker caste. (2) We suggest that maternal certainty of offspring dispersal decisions is one of the main constraints on the evolution of a morphological worker caste in the facultatively eusocial and cooperatively breeding Hymenoptera. (3) The decoupling of control over morphology and behaviour could be a key factor maintaining a morphological worker caste, and can lead to reduced conflict between offspring and provisioners. (4) The productivity of workers in the absence of specialist worker morphology can affect the likelihood of specialist morphology being favoured by selection.

## Acknowledgements

This work was funded by the University of Exeter. We would like to thank Alex Wild for kindly allowing use of the photograph for Figure 1. We would also like to thank Andy Gardner, Peter Nonacs, and two anonymous reviewers for their comments on the manuscript, and Vincent Jansen for helpful discussions about the models.

## Data Availability Statement

This study did not generate any new data. Additional information supporting this publication is available as supplementary information accompanying this publication.

