## Supplementary Material for "The evolution of morphological castes under decoupled control"

### ***Part 1 Mother and daughter preference for workers***

Behavioural castes are often mutable, so that individuals can switch castes over their lifetimes. The inclusion of effects such as worker takeover, through which a worker can become a reproductive, may reduce the threshold worker productivity required for a daughter to favour joining a behavioural worker caste (Field et al., 1999; Toyoizumi and Field, 2014; Price and Field, 2022). Similarly, if a daughter can help raise more reproductive sisters (r=0.75) than reproductive brothers (r=0.25) by staying as a worker, it will further lower the productivity of a worker relative to a reproductive required for the daughter to favour joining a behavioural worker caste (Trivers and Hare, 1976). The fact that we see evidence for behavioural worker castes preceding morphological worker castes in the social insects (Bourke, 2011; and see main text Discussion) therefore indicates that at least one of the following must be true: (1) the mutability of behavioural worker castes causes daughters to favour joining them at a lower worker productivity than required for mothers to favour imposing a morphological worker caste, (2) workers help raise broods which are sufficiently female biased to mean that daughters favour joining behavioural worker castes at a lower worker productivity than required for mothers to favour imposing a morphological worker caste or (3) the evolution of a morphological worker caste is reliant on the prior presence of a behavioural worker caste.

If we let $x$ be the number of offspring of relatedness 0.5 a solitary individual produces, $q$ the productivity of a replacement queen relative to a solitary individual, $p$ the probability of a worker taking over the queen position (at any point in its life), $A$ the average age of an inheriting worker (from 0-1, as a proportion of worker lifespan), and $h$ the relative productivity of a worker to a solitary individual, then we find the following.

The fitness of a daughter working when worker takeover is possible, relative to a reproductive daughter, can be broken down into three components; the fitness an individual would receive from working if it never inherits ($(1-p)hx$), the fitness an individual would receive from working if it later inherits ($Aphx$), and the fitness an individual would receive once it inherits ($(1-A)qxp$). We can then generate the following:

$$F_{dwt} = (1-A)qxp + (1-p)hx + Aphx = (1-A)qxp + [1+p(A-1)]hx$$

The fitness of a daughter dispersing as a solitary individual reproductive:

$$F_{dr} = x$$

The fitness a mother receives from a daughter working when takeover is possible:

$$F_{mwt} = \frac{(1-A)qxp}{2} + [1+p(A-1)]hx$$

The fitness a mother receives when a daughter disperses as a solitary individual:

$$F_{mr} = \frac{x}{2}$$

The solution for equal mother and daughter preference for worker daughters is therefore when both, (1) working gives a daughter higher fitness than dispersing as a reproductive;

$$\frac{F_{dwt}}{F_{dr}} > 1$$

Which simplifies to:

$$(1 - A)qp \ + [1 + p(A - 1)]h > 1$$

And (2) mothers get the same proportional increase to fitness from daughters working instead of dispersing as daughters do;

$$\frac{F_{mwt}}{F_{mr}} = \frac{F_{dwt}}{F_{dr}}$$

or

$$\frac{F_{mwt}}{F_{mr}} \div \frac{F_{dwt}}{F_{dr}} = 1$$

Which simplifies to:

$$\frac{(1 - A)qp \ + \ 2[1 + p(A - 1)]h}{(1 - A)qp \ + [1 + p(A - 1)]h} = 1$$

And therefore requires at least one of the following be true:

$h = 0$ , $q = \infty$, or both $p = 1$ and $A = 0$.

As such, a mother and a daughter will both prefer daughters to become helpers instead of reproductives at the same worker productivity (so both (1) and (2) are true) when either;

1.) $h = 0$ and $(1 - A)qp > 1$ (worker daughters do not actually work, but have a higher expected fitness from staying with the expectation of inheriting than from founding their own nest)

2.) $p = 1$, $A = 0$, and $q > 1$ (worker daughters always immediately inherit the queen position, and are more productive than solitary foundresses when they do)

3.) $q = \infty$ and $p > 0$ (worker daughters inheriting the queen position are infinitely productive and worker daughters do sometimes inherit)

On its own, the mutability of behavioural castes (explanation 1) therefore cannot reasonably affect the argument that mothers will impose a worker caste before daughters prefer to join it. This would require workers to always immediately inherit the queen position or to produce an infinite number of offspring whenever they do.

Split sex ratios (Seger, 1983; Grafen, 1986; Godfray and Grafen, 1988; Godfray, 1990; Gruber and Field, 2022) are also unlikely to result in brood sex ratios that are female-biased enough to significantly impact this argument (Godfray and Hardy, 1990; Grafen, 1986; Boomsma and Eickwort, 1993). But even when daughters raise entirely sisters, they receive only 1.5 times the fitness benefit of raising the same number of their own offspring, less than the doubled benefit that mothers receive.

To see how these effects (explanations 1 and 2) work in tandem, with worker daughters helping to raise only sisters, then we can instead say;

The fitness of a daughter working when worker takeover is possible is:

$$F_{dwt} = (1 - A)qxp \; + \frac{3[1 + p(A - 1)]hx}{2}$$

While everything else remains the same as before.

The solution for equal mother and daughter preference for worker daughters then becomes when;

$$(1 - A)qp \; + \frac{3[1 + p(A - 1)]h}{2} > 1$$

And

$$\frac{(1 - A)qp \; + \; 2[1 + p(A - 1)]h}{(1 - A)qp \; + \frac{3[1 + p(A - 1)]h}{2}} = 1$$

Since $2 > \frac{3}{2}$, our previous results remain true, and worker daughters helping to raise all female (sister) broods cannot enable mothers and daughters to both prefer daughters to become helpers instead of reproductives at the same worker productivity. Thus, we find that our proposed explanations 1 and 2, even acting in tandem, could not cause daughters to favour the arisal of a worker caste before mothers do.

### ***Part 2 Generalised model and mutable behavioural castes***

We can treat selection on morphology more generally by considering two individuals, **A** and **B**, where individual **A** controls the morphology of individual **B** but individual **B** retains full control over its own behaviour (for traits which individuals control their own expression of, e.g. under recoupled control, individual **A** and individual **B** are then the same individual). Individual **B** will express one of two behavioural types ($\boldsymbol{x}$, denoted type 1 and type 2) and, if we wish to include mutability of behaviour, may change which of these two types it expresses over its lifetime. If we include mutability of behaviour, we must make two additional assumptions, which are that the lifespan of individual **B** is independent of its behavioural type, and fitness ($\boldsymbol{v_x}$ or $\boldsymbol{v'_x}$ ; Table S1) is independent of the age of individual **B**. It is worth noting that in most systems, these two additional assumptions will not hold, but they should be adequate for our purposes in the following text. Our parameters under these assumptions are detailed in Table S1. Perhaps most important of these is our parameter $\boldsymbol{u}$ which represents the expected proportion of time spent as behavioural type 1 by individual **B**, enabling us to relax our previous assumption of behavioural caste immutability. For instance, if we want to consider the fitness a mother stands to gain by imparting a morphological trait onto a daughter in B1, as in our examples in Equations 1-6, assuming now that daughters which stay as workers have a chance of inheriting the role of queen from their mother, we can generate a value for $\boldsymbol{u}$ as follows: if we set the behavioural worker caste as behavioural type 2 and the reproductive caste as behavioural type 1, with a 40% chance of a worker inheriting ($\boldsymbol{P.inherit}$), an average time of inheritance of half way through worker lifespan ($\boldsymbol{T.inherit}$), and probability of dispersing of 0.4 ($\boldsymbol{d}$), we can calculate a value of $\boldsymbol{u}$ =0.52, where $\boldsymbol{u} = \boldsymbol{d} + (\boldsymbol{1} - \boldsymbol{d}) * [\boldsymbol{P.inherit} * \boldsymbol{T.inherit}]$.

*Table S1. Parameters used in Equation S1. Here we assume $\boldsymbol{m}$ is a trait which individual* ***B*** *can respond to.*

| | With morphological trait $\boldsymbol{m}$ | Without morphological trait $\boldsymbol{m}$ ("baseline value") |
|---|---|---|
| Fitness individual **B** will provide to individual **A** over its lifetime if it only ever expresses behavioural type 1 | $\boldsymbol{v_1'}$ | $\boldsymbol{v_1}$ |
| Fitness individual **B** will provide to individual **A** over its lifetime if it only ever expresses behavioural type 2 | $\boldsymbol{v_2'}$ | $\boldsymbol{v_2}$ |
| Cost to individual **A** represented by individual **B** | $\boldsymbol{c'}$ | $\boldsymbol{c}$ |
| Expected proportion of time spent as behavioural type 1 by individual **B** when trait $\boldsymbol{m}$ increases the average proportion of time spent as behavioural type 2 | $\boldsymbol{u(1-z)}$ | $\boldsymbol{u}$ |
| Expected proportion of time spent as behavioural type 1 by individual **B** when trait $\boldsymbol{m}$ increases the average proportion of time spent as behavioural type 1 | $\boldsymbol{1-(1-u)(1+z)}$ | $\boldsymbol{u}$ |
| Expected fitness per unit investment individual **B** will provide to individual **A** | $\boldsymbol{f'}$ | $\boldsymbol{f}$ |

Using the parameters in Table S1 we can therefore generate Equation S1, which is represented in Figure S1B:

***Equation S1:***

$$\boldsymbol{f = \frac{uv_1 + (1-u)v_2}{c}},$$

$$\boldsymbol{f' = \begin{cases} \dfrac{u(1-z)v_1' + [1-u(1-z)]v_2'}{c'}, & z \in \{0 \leq z \leq 1\} \\[2ex] \dfrac{[1-(1-u)(1+z)]v_1' + (1-u)(1+z)v_2'}{c'}, & z \in \{-1 \leq z \leq 0\} \end{cases}}$$

Which we see shares the same functional form as equation 4 in the main text. It is worth noting that if offspring are only altering their dispersal choices, not their expected likelihood or time of inheriting, as a response to the presence of a morphological trait then it may be more informative to substitute $\boldsymbol{a}$ and $\boldsymbol{d}$ into Equation S1, where;

$\boldsymbol{u(1-z) = d(1-a) + [1 - d(1-a)](P.inherit * T.inherit)}$, when $\boldsymbol{a > 0}$

and

$\boldsymbol{(1-u)(1+z) = (1-d)(1+a)(1 - [P.inherit * T.inherit])}$, when $\boldsymbol{a < 0}$.

The relationship between $\boldsymbol{a}$ and $\boldsymbol{z}$ can be most clearly seen illustrated in Figure S2. Here we see illustrated how $\boldsymbol{a}$ modifies the likelihood of dispersal but not of inheritance, while $\boldsymbol{z}$ is inclusive of both changes to dispersal and inheritance.

It is also worth mentioning that specialist worker morphology, which decreases the fitness per unit cost of a reproductive daughter, can now be selected for even when all offspring become reproductives ($\boldsymbol{u = 1}$) if offspring with trait *m* are more likely to stay ($\boldsymbol{a} > 0$; Figure 2, Figure S1). However, it seems very unlikely that this combination will exist: daughters are unlikely to have evolved the ability to respond to the presence of trait $\boldsymbol{m}$ in a population where no daughters stay as workers in its absence. The benefit (per unit cost) in fitness from a worker would also have to be very large relative to the loss in fitness (per unit cost) from a reproductive (e.g. a very favourable trait) to enable this. As such we consider that our conclusion from the main text, that specialist worker morphology cannot evolve without the prior evolution of helping behaviour, remains true.

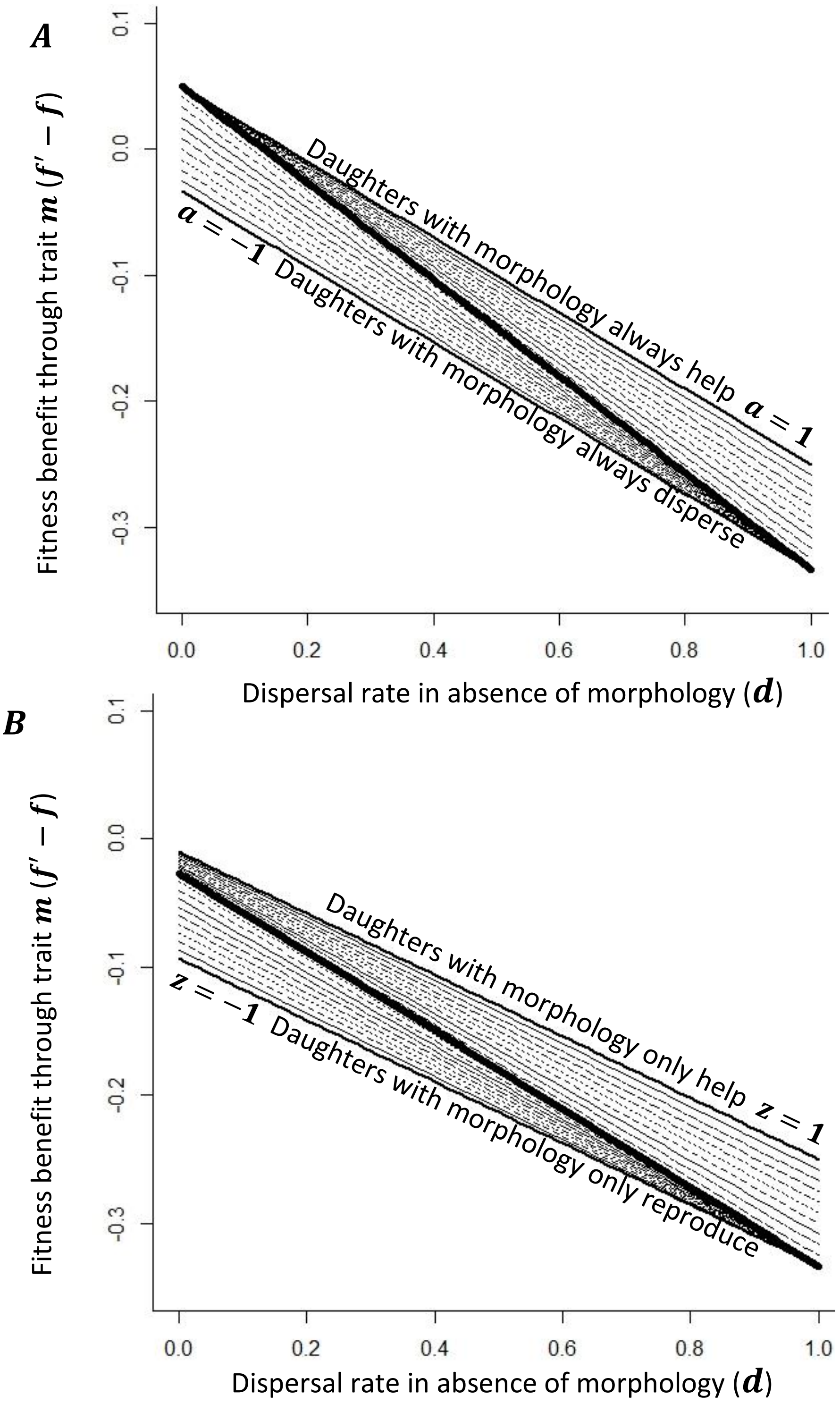


Figure S1. The effect of worker takeover on the relationship between the net fitness increase a provisioner will receive ($f' - f$) from imparting morphological trait $m$ on to a daughter, and the baseline probability of a daughter dispersing. Plotted for $-1 \leq a \leq 1$ and $-1 \leq z \leq 1$ as appropriate at 0.1 increments. The thick central lines represent $a = 0$ and $z = 0$. $A$, $v'_w > v'_r$ for parameter values ($v'_w$ = 0.9, $v'_r$ = 0.8, $v_w$ = 0.7, $v_r$ = 1, $c'$ = 1, $c$ = 1). $B$, uses the same parameter values as panel A, however we have assumed worker takeover with a 40% chance of a worker inheriting ($P.inherit$) and average time of inheritance of half way through worker lifespan ($T.inherit$). $u$ has been calculated such that $u = d + (1 - d) * [P.inherit * T.inherit]$, as given in supplementary materials part 2.

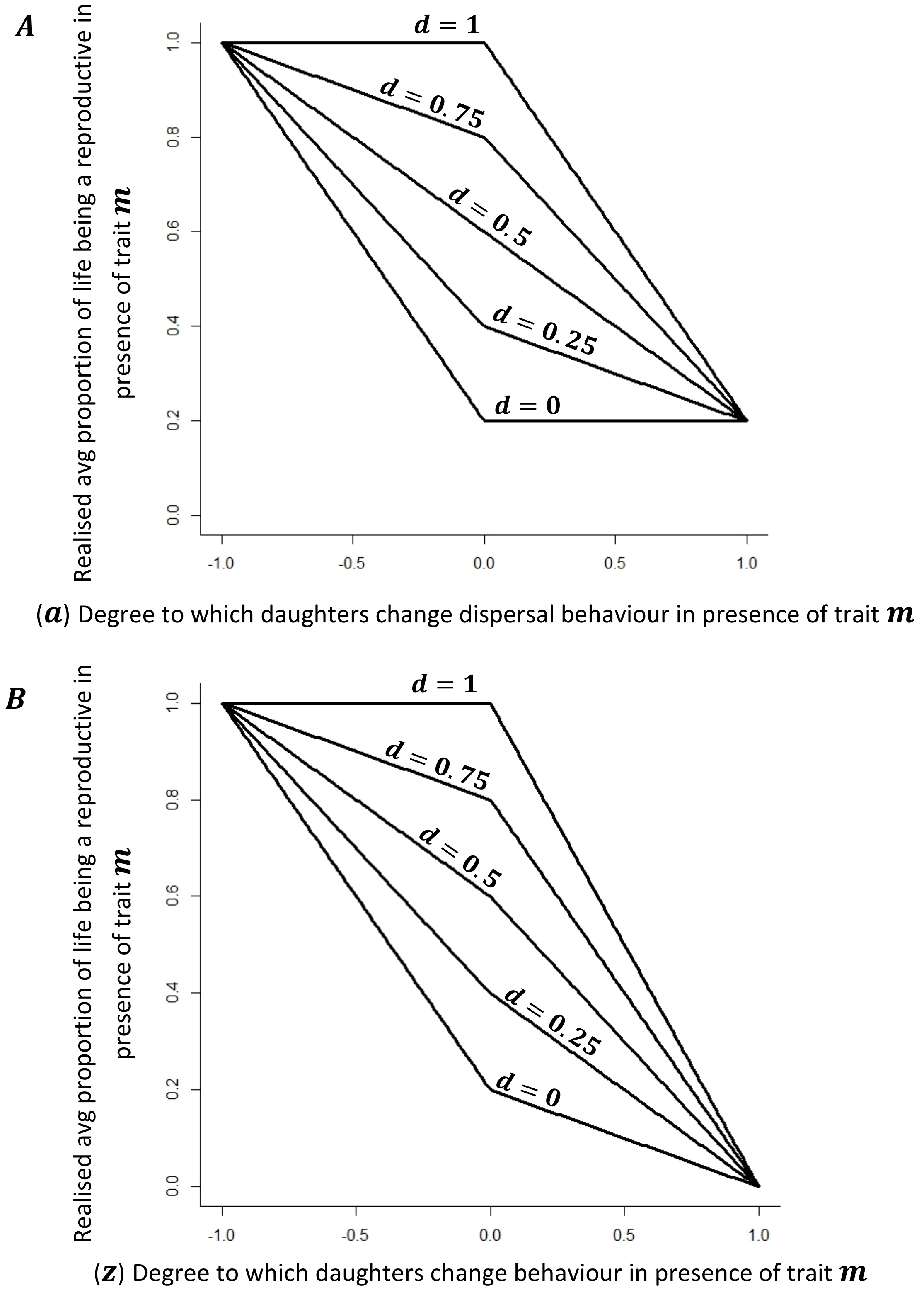


*Figure S2. The relationship between* $\boldsymbol{a}$ *(the degree to which daughters change their dispersal behaviour in the presence of trait* $\boldsymbol{m}$*),* $\boldsymbol{z}$ *(the degree to which daughters change their likelihood of being a reproductive in the presence of trait* $\boldsymbol{m}$*), and realised average proportion of life an offspring spends as a reproductive. Different lines represent different values of baseline dispersal (dispersal in the absence of trait* $\boldsymbol{m}$*;* $\boldsymbol{0 \leq d \leq 1}$*) in 0.25 increments. For parameter values (*$\boldsymbol{P.inherit}$*= 0.4,* $\boldsymbol{T.inherit}$ *=0.5).* $\boldsymbol{A}$ *varying* $\boldsymbol{a}$ *(the degree to which daughters change their dispersal behaviour in the presence of trait* $\boldsymbol{m}$*).* $\boldsymbol{B}$*, varying* $\boldsymbol{z}$ *(the degree to which daughters change their likelihood of being a reproductive in the presence of trait* $\boldsymbol{m}$*).*

#### *Part 3 Trait benefit as independent of baseline productivity*

Further to the linear relationship discussed in the paper (Equation 6), we now consider a second case: where the fitness benefit provided by trait $\boldsymbol{m}$ is independent of $\boldsymbol{v_w}$ and equal to the constant $\boldsymbol{k}$, such that $\boldsymbol{v'_w = v_w + k}$. An example would be when each large worker increases the reproductive output of a nest by one more offspring than each small worker. Since the difference in productivity between small and large workers is independent of how productive small workers are, we might intuitively expect the likelihood of trait $\boldsymbol{m}$ being favoured by selection to be independent of $\boldsymbol{v_w}$.

Substituting $\boldsymbol{v'_w = v_w + k}$ into Equation 5, we obtain:

*Equation S2:*

$$\boldsymbol{0 < \begin{cases} v_w(d[c' + (a-1)c] + c - c') + [1 - d(1-a)]kc + d[cv'_r(1-a) - c'v_r], & a \in \{0 \le a \le 1\} \\ v_w(d[c' - (1+a)c] + c(1+a) - c') + (1-d)(1+a)kc + [1 - (1-d)(1+a)]cv'_r - dc'v_r, & a \in \{-1 \le a \le 0\} \end{cases}}$$

When $\boldsymbol{a = 0}$ and $\boldsymbol{c = c'}$ we see our prediction is true, and the likelihood that trait $\boldsymbol{m}$ is favoured by selection is independent of $\boldsymbol{v_w}$ (Figure S3). However, if offspring change their behaviour in the presence of trait $\boldsymbol{m}$ ($\boldsymbol{a \neq 0}$, and $\boldsymbol{0 < d < 1}$; Figure 3), the likelihood that trait $\boldsymbol{m}$ is favoured is now correlated with the fitness of workers lacking trait $\boldsymbol{m}$ ($\boldsymbol{v_w}$; Figure S3). This is because trait $\boldsymbol{m}$ not only changes the productivity of daughters, as both reproductives and workers, but also the likelihood of a mismatch occurring when a daughter has trait $\boldsymbol{m}$. When $\boldsymbol{a > 0}$ (and $\boldsymbol{d > 0}$; Figure 3) we see a positive relationship between $\boldsymbol{v_w}$ and likelihood of trait $\boldsymbol{m}$ evolving. Since $\boldsymbol{a > 0}$, the presence of trait $\boldsymbol{m}$ is increasing the occurrence of worker behaviour. With $\boldsymbol{v'_w = v_w + k}$, as $\boldsymbol{v_w}$ increases, the productivity of workers with trait $\boldsymbol{m}$ also increases. These two effects together give us the resultant positive relationship between $\boldsymbol{v_w}$ and likelihood of trait $\boldsymbol{m}$ evolving (Figure S3); an increased likelihood of working in the presence of trait $\boldsymbol{m}$ will give greater fitness returns as the productivity of workers with trait $\boldsymbol{m}$ increases.

When $\boldsymbol{v_w} > \boldsymbol{v'_r} - \boldsymbol{k}$, this increased likelihood of worker behaviour in the presence of trait $\boldsymbol{m}$ ($\boldsymbol{a} > \boldsymbol{0}$) is then increasing the likelihood of trait $\boldsymbol{m}$ evolving (Figure S3). This is because $\boldsymbol{v_w} = \boldsymbol{v'_r} - \boldsymbol{k}$ represents the point beyond which $\boldsymbol{v'_w} > \boldsymbol{v'_r}$ (since $\boldsymbol{v'_w} = \boldsymbol{v_w} + \boldsymbol{k}$), such that an offspring with trait $\boldsymbol{m}$ working results in a higher fitness than that offspring dispersing, and hence the point beyond which an increased likelihood of worker behaviour in the presence of trait $\boldsymbol{m}$ ($\boldsymbol{a} > \boldsymbol{0}$) provides a selective advantage.

This relationship is reversed when $\boldsymbol{a} < \boldsymbol{0}$, and we would then see the likelihood of trait $\boldsymbol{m}$ evolving decrease with $\boldsymbol{v_w}$. This effect of $\boldsymbol{a}$ on mismatch is what enables the previously observed ability for very small values of $\boldsymbol{a}$ ($\boldsymbol{a} < -\boldsymbol{0.5}$, so a strong behavioural bias towards dispersal) to overturn the positive relationship between $\boldsymbol{v_w}$ and the likelihood of trait $\boldsymbol{m}$ being selected seen in Equation 6 (Figure 4). However, since we expect specialist worker morphology to favour worker daughters more than reproductive daughters ($\boldsymbol{k} > \boldsymbol{j}$, where $\boldsymbol{v'_r} = \boldsymbol{v_r} + \boldsymbol{j}$), we would expect $\boldsymbol{v'_w} > \boldsymbol{v'_r}$ to be satisfied at relatively small values of $\boldsymbol{v_w}$. Interestingly, while workers with the trait provide less fitness than reproductives with the trait ($\boldsymbol{v'_w} < \boldsymbol{v'_r}$), even specialist morphological traits, which increase the productivity of workers and decrease that of reproductives, will benefit mothers most when offspring disperse more frequently ($\boldsymbol{a} < \boldsymbol{0}$) (Figure 4, Figure S3, Figure S4, Figure S5)**.** It is also the case that offspring would only be expected to disperse more frequently in the presence of a trait ($\boldsymbol{a} < \boldsymbol{0}$) when workers with the trait expect to be less productive than reproductives with the trait ($\boldsymbol{2v'_w} < \boldsymbol{v'_r}$; under standard assumptions of: equal sex allocation, no worker takeover and singly mated queens with full reproductive control). However, as we establish in the main text, with no morphological specialisation we would tend to expect working to occur only when daughters are at least as productive as workers as when they are reproductives (standard assumptions; Figure 5). If workers are sufficiently morphologically specialised, we would instead expect provisioners to control when offspring disperse (Figure 5). This means that we could only ever expect working to occur when a provisioner gets more fitness from an offspring which works than from an offspring which disperses, which is when $\boldsymbol{v_w} > \boldsymbol{v_r}$ (under standard assumptions this would be when workers are half as productive as reproductives). Since $\boldsymbol{v'_w} - \boldsymbol{v_w} > \boldsymbol{v'_r} - \boldsymbol{v_r}$ is by definition true for a specialist worker trait, we can then see that $\boldsymbol{v'_w} > \boldsymbol{v'_r}$ and $\boldsymbol{a} \geq \boldsymbol{0}$ represents the parameter space we are most interested in for the evolution of specialist worker morphological traits in the social Hymenoptera.

As Fisher first highlighted in 1930, it is the fitness returns per unit investment which principally guides evolutionary outcomes, not just fitness alone. We can see this reflected in Equation S2 when $\boldsymbol{c} \neq \boldsymbol{c'}$ (the cost represented by a daughter with trait $\boldsymbol{m}$ differs from the cost without it): a relationship emerges between the likelihood of trait $\boldsymbol{m}$ evolving and $\boldsymbol{v_w}$ (Figure S4), despite the extra benefit of possessing trait $\boldsymbol{m}$ ($\boldsymbol{k}$) being independent of baseline productivity (and the fitness a mother gets from this baseline productivity $\boldsymbol{v_w}$). When $\boldsymbol{c} > \boldsymbol{c'}$ this relationship is positive - trait $\boldsymbol{m}$ is more likely to be favoured when baseline fitness is larger - and when $\boldsymbol{c} < \boldsymbol{c'}$ the relationship is negative (Figure S4). This is because when $\boldsymbol{c} \neq \boldsymbol{c'}$ we are changing the cost of producing a daughter with trait $\boldsymbol{m}$: so while the benefit of trait $\boldsymbol{m}$ ($\boldsymbol{k}$) remains constant, as the baseline productivity (and hence $\boldsymbol{v_w}$) increases, the benefit per unit cost does not. We can illustrate this by considering the following; for a trait $\boldsymbol{m}$ to confer a net benefit per unit investment to the fitness represented by a worker daughter, the per unit cost baseline fitness from a worker must be less than the per unit cost fitness from a worker with the trait ($\frac{v_w}{c} < \frac{v_w+k}{c'}$ must be true). When $k$ is constant and independent of $v_w$, as our value of $v_w$ increases the benefit of trait $m$ ($k$) will become less important in determining the outcome of $\frac{v_w}{c} < \frac{v_w+k}{c'}$ than the difference between $c$ and $c'$. For instance, if $c' = 2c$, then the benefit per unit cost of a trait $m$ will be $\frac{v_w+k}{2c} - \frac{v_w}{c}$, which is greatest when $v_w$ is small (e.g. when $k = 2$, we get a benefit per unit cost of $\frac{2}{c}$ when $v_w = 2$, but a benefit per unit cost of $-\frac{4}{c}$ when $v_w = 10$). More generally we can say when $k$ is constant and $c < c'$, $\frac{v_w}{c} < \frac{v_w+k}{c'}$ is most likely to be true when $v_w$ is small, while if $c > c'$, $\frac{v_w}{c} < \frac{v_w+k}{c'}$ is most likely to be true when $v_w$ is large. Therefore the costlier a specialist worker trait is, the more likely it is to evolve when worker productivity is low (Figure S4A), while the cheaper a specialist worker trait is, the more likely it is to evolve when worker productivity is high (Figure S4B). Returning to our earlier example in Equation 6, where $v'_w = 2v_w$, $\frac{v_w}{c} < \frac{2v_w}{c'}$ must then be true for trait $m$ to be favoured by selection (without favourably biasing mismatch; Figure 4, Figure S4). It then follows that If $2c < c'$, $\frac{v_w}{c} < \frac{2v_w}{c'}$ is closest to being true when $v_w$ is small (though cannot be fulfilled without the addition of a constant $k$ to give $\frac{v_w}{c} < \frac{2v_w+k}{c'}$), while if $2c > c'$, $\frac{v_w}{c} < \frac{2v_w}{c'}$ is always true but greatest when $v_w$ is large. It thus follows that, more generally, if a morphological trait is sufficiently costly it will be most likely to evolve when $v_w$ is small, otherwise that trait will be most likely to evolve when $v_w$ is large. However, the degree to which the cost of offspring with the trait ($c'$) and without it ($c$) must differ for this to be the case will vary as a function of the relationship between $v'_w$ and $v_w$. Importantly though, we always predict a cheaper morphological trait to evolve over a more expensive trait which provides the same benefit, regardless of our value of $v_w$ (Figure S4).

Note: for our treatments in this section we have assumed that for a given trait $\frac{\mathrm{d}v'_w}{\mathrm{d}v_w}$ is either always greater or less than $\frac{c'}{c}$, and never crosses this value (this can also be expressed in terms of $\frac{\mathrm{d}(v'_w - v_w)}{\mathrm{d}v_w}$ relative to $\frac{c'-c}{c}$). We believe this to be appropriate for the majority of envisionable cases. But should this not be the case, the extraction of more general predictions, as we have attempted here, becomes difficult.

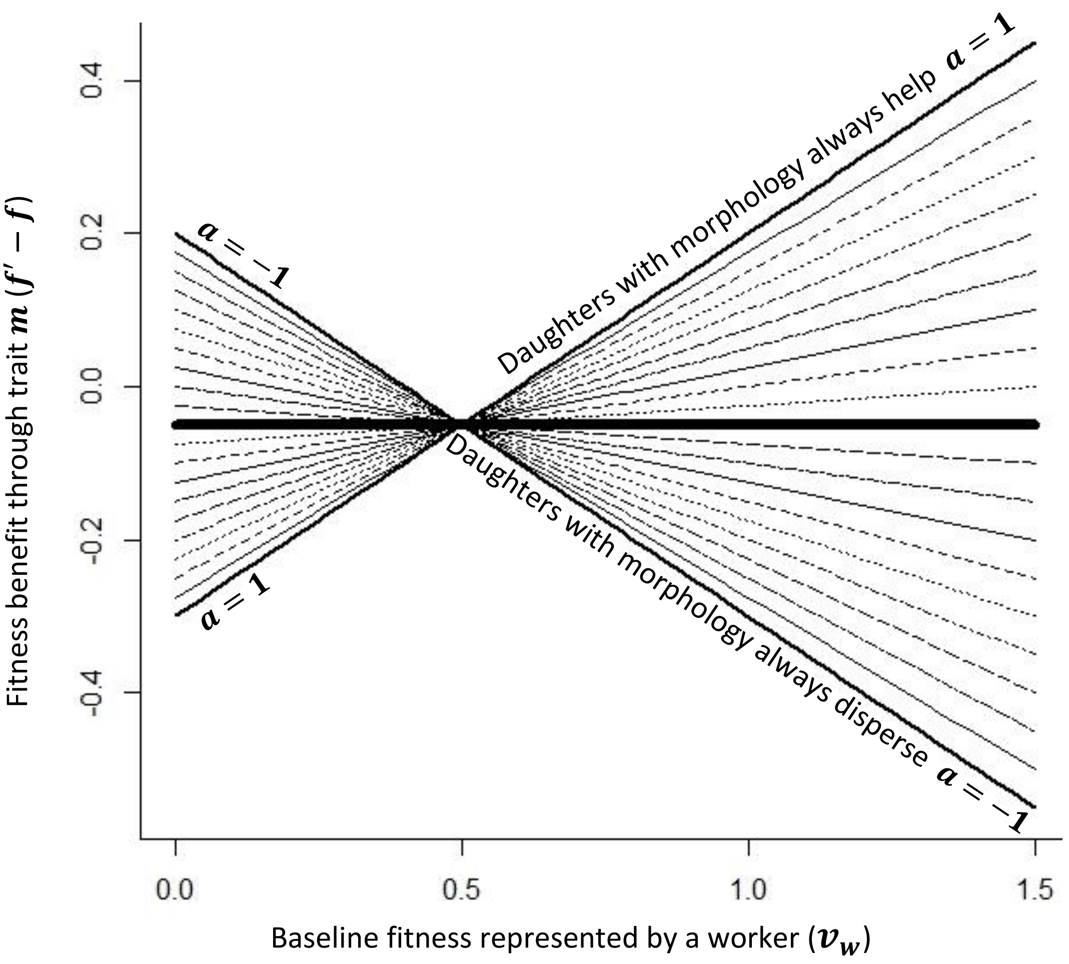


Figure S3. The ability for offspring to respond to morphology mediates the relationship between the baseline fitness (or productivity) of a worker and the likelihood of a specialist morphological trait $m$ being favoured by selection. We have plotted this for $-1 \leq a \leq 1$ at 0.1 increments using parameter values ($k = 0.2$, $v'_r = 0.7$, $v_r = 1$, $d = 0.5$, $c' = 1$, $c = 1$), meaning trait $m$ favours workers. $v_w = v'_r - k$ when $v_w = 0.5$, which is just half our value of $v_r$. The thick central line represents $a = 0$.

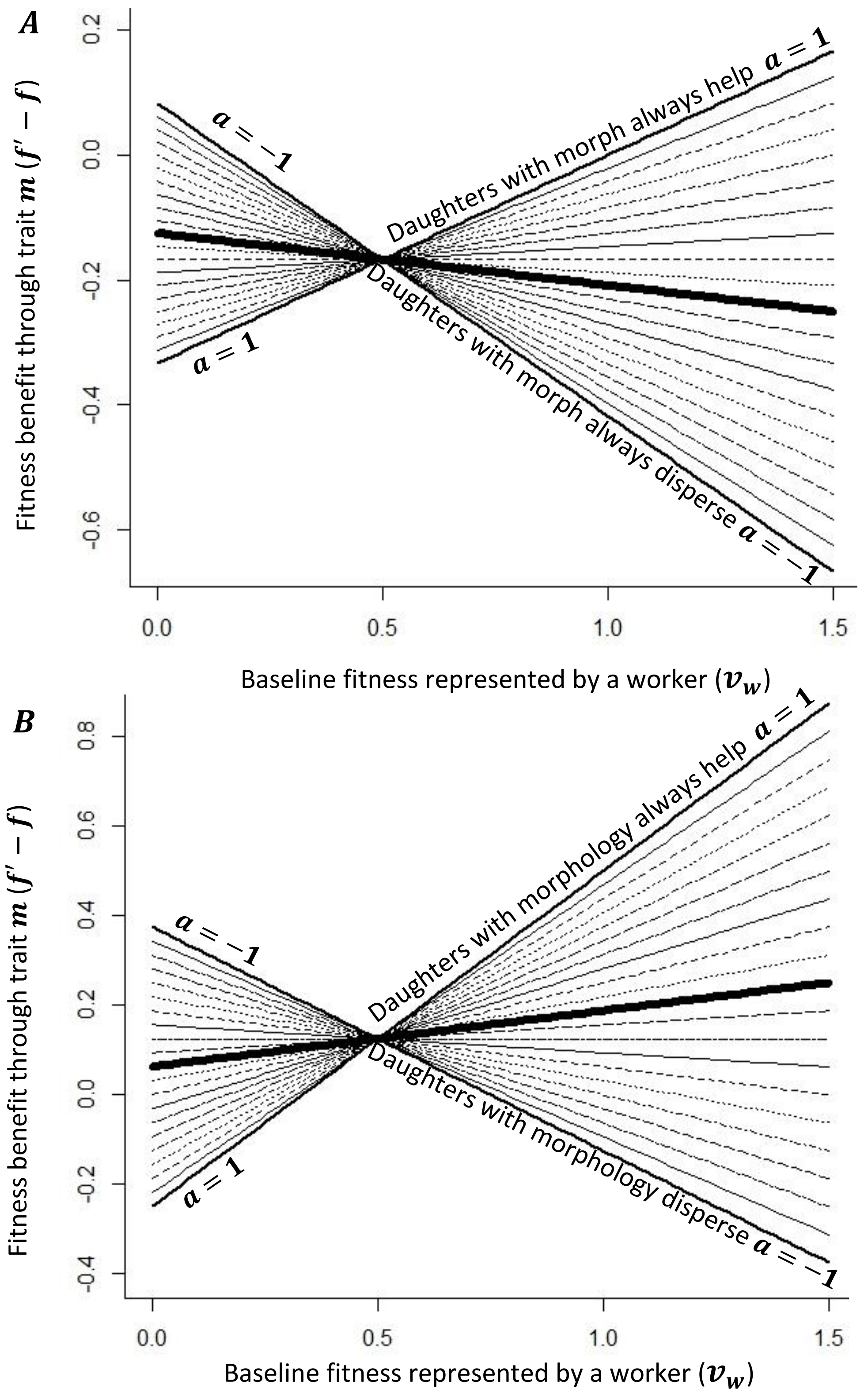


*Figure S4. When $c \neq c'$ there is a linear relationship between the net fitness increase a provisioner will receive ($f' - f$) from imparting morphological trait $m$ on to a daughter, and the baseline fitness (or productivity) of a worker. We have plotted this for $-1 \leq a \leq 1$ at 0.1 increments. The thick central lines represent $a = 0$. $A$, $c' > c$ ($k$ = 0.2, $v_r'$ = 0.7, $v_r$ = 1, $d$ = 0.5, $c'$ = 1.2, $c$ = 1). $B$, $c' < c$ ($k$ = 0.2, $v_r'$ = 0.7, $v_r$ = 1, $d$ = 0.5, $c'$ = 0.8, $c$ = 1).*

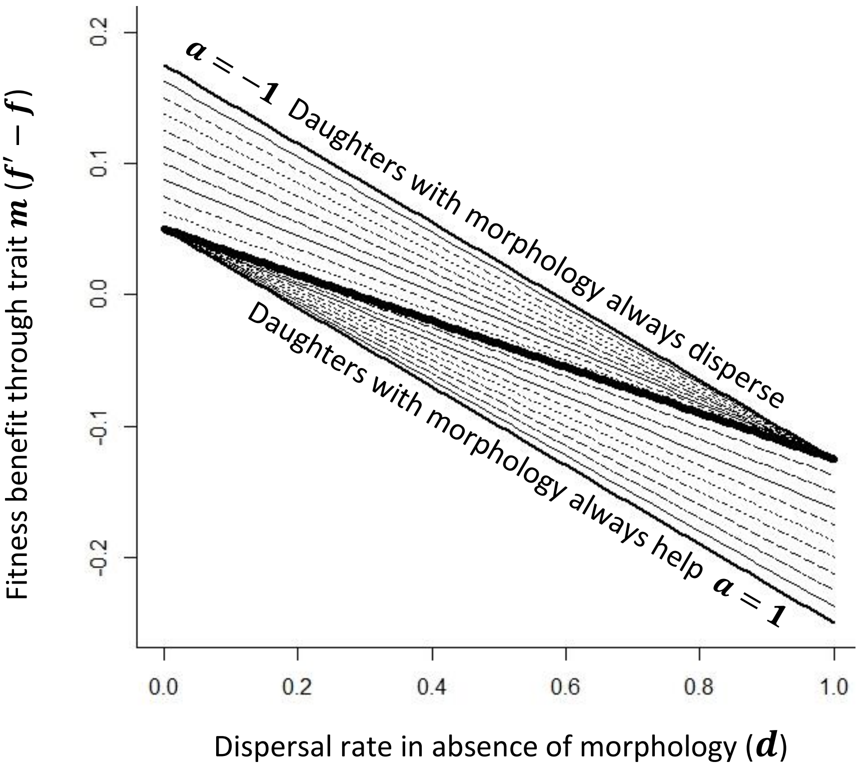


*Figure S5. Paired with Figure 2, shows the linear relationship between the net fitness increase a provisioner will receive from imparting morphological trait $\boldsymbol{m}$ onto a daughter ($\boldsymbol{f' - f}$), and the baseline probability of a daughter dispersing (see Table 1 for parameter definitions). We have plotted this for $-\mathbf{1} \le \boldsymbol{a} \le \mathbf{1}$ at 0.1 increments. The thick central lines represent $\boldsymbol{a} = \mathbf{0}$ and hence Equation 3. Here daughters with trait $\boldsymbol{m}$ give less fitness as workers than reproductives ($\boldsymbol{v'_w} < \boldsymbol{v'_r}$) for parameter values ($\boldsymbol{v'_w}$ = 0.6, $\boldsymbol{v'_r}$ = 0.7, $\boldsymbol{v_w}$ = 0.7, $\boldsymbol{v_r}$ = 1, $\boldsymbol{c'}$ = 0.8, $\boldsymbol{c}$ = 1).*

***Part 4          Bivoltine cooperatively breeding haploid model***

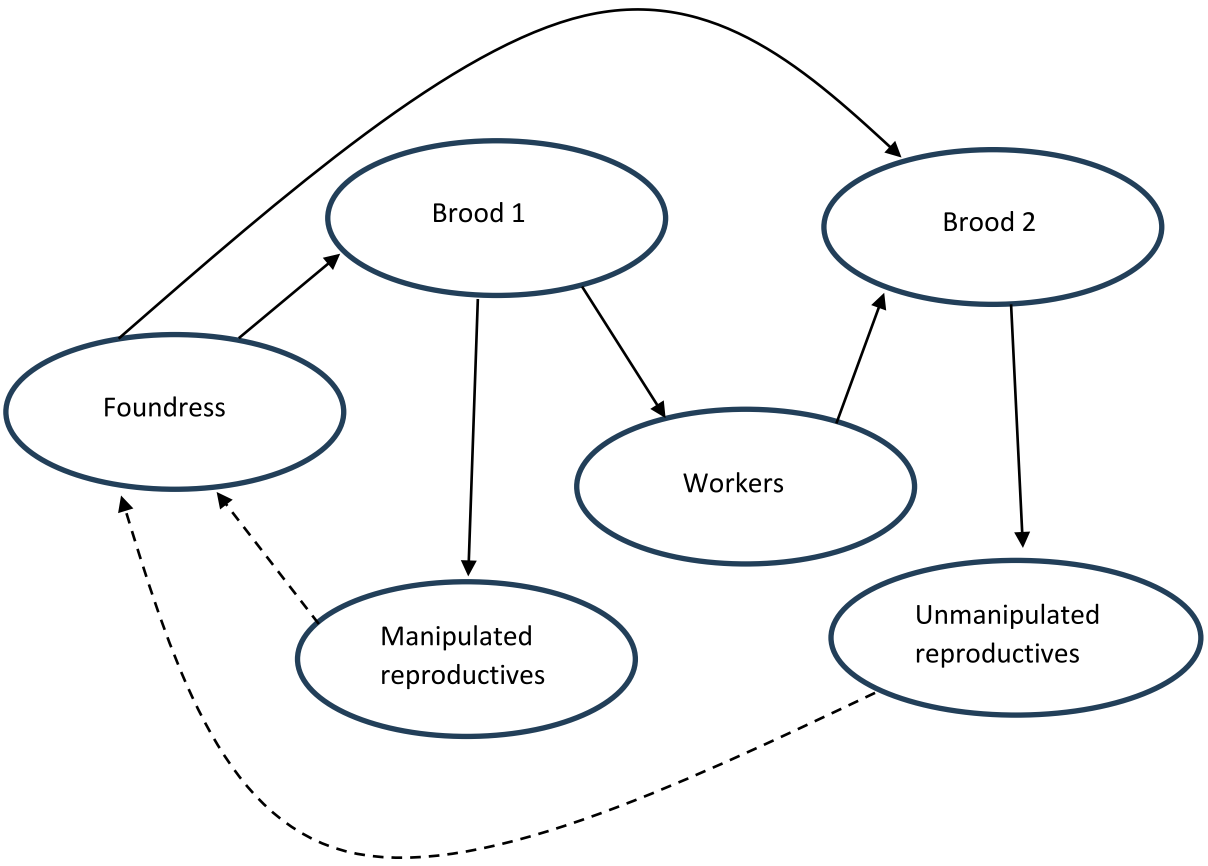


*Figure S6. Imagined seasonal life cycle of our bivoltine cooperatively breeding haploid system. Dotted lines represent the transition from one season to the next.*

**Parameters**

$\boldsymbol{g_x}$ = Breeding value (genetic component of phenotype) of a foundress with genotype $x$

$\boldsymbol{\Lambda(g_x)}$ = Investment of a worker into brood 2, as a function of the foundress's breeding value

$\boldsymbol{\gamma_u}$ = Investment of an unmanipulated foundress into a brood

$\boldsymbol{\gamma_m(g_x)}$ = Investment of a manipulated foundress into a brood, as a function of its breeding value

$\boldsymbol{z(g_x)}$ = Founding competitiveness of a manipulated reproductive relative to an unmanipulated reproductive, as a function of the foundress's breeding value

$\boldsymbol{c_u}$ = Cost of producing an unmanipulated offspring

$\boldsymbol{c_m(g_x)}$ = Cost of producing a manipulated offspring, as a function of the foundress's breeding value.

$\boldsymbol{d}$ = Dispersal rate of offspring produced in brood 1

$\boldsymbol{m_x}$ = Number of manipulated reproductives of genotype $x$ present at the start of a season

$\boldsymbol{u_x}$ = Number of unmanipulated reproductives of genotype $x$ present at the start of a season

$\boldsymbol{N_x} = u_x + m_x$ =Total number of reproductives of genotype $x$ present at the start of a season

$\boldsymbol{N'_x} = u'_x + m'_x$ =Total number of reproductives of genotype $x$ present at the end of a season

$\boldsymbol{n}$ = Number of nest sites in the population

$\boldsymbol{x}$ = Genotype, $x = 1$ for the resident genotype and $x = 2$ for the mutant phenotype

### Model derivations

We imagine a bivoltine cooperatively breeding Hymenopteran with a life cycle as given in Figure S6. We assume a fixed number of nest sites which the reproductives, produced in the previous season, must compete for to found a nest at the start of a season. If a reproductive successfully founds a nest it becomes a foundress. Foundresses produce two broods over a season, which they invest equally into. The first brood they may manipulate morphologically, and a proportion of the offspring produced will disperse as (manipulated) reproductives, while the remainder stay as workers. For our first model, we assume the proportion of brood 1 which stay as helpers does not vary as a function of manipulation. The second brood is not manipulated, and all offspring produced from it disperse. The second brood also benefits from the contributions of any offspring from the first brood which stayed as helpers.

*Model 1 (fixed dispersal)*

Assuming an infinite number of nest sites $n$, we can then write the following;

***Equation S3***

$$m'_x = \frac{n[u_x\gamma_u \ + \ m_x z(g_x)\gamma_m(g_x)]}{u_1 + \ u_2 + z(g_1)m_1 + z(g_2)m_2} \ [\frac{d}{c_m(g_x)}]$$

***Equation S4***

$$u'_x = \frac{n[u_x\gamma_u \ + \ m_x z(g_x)\gamma_m(g_x)]}{u_1 + \ u_2 + z(g_1)m_1 + z(g_2)m_2} \ [\frac{1}{c_m(g_x)}(1-d)\Lambda(g_x) + \frac{1}{c_u}]$$

***Equation S5***

$$N'_x = \frac{n[u_x\gamma_u \ + \ m_x z(g_x)\gamma_m(g_x)]}{u_1 + \ u_2 + z(g_1)m_1 + z(g_2)m_2} \ [\frac{1}{c_m(g_x)}(1-d)\Lambda(g_x) + \frac{1}{c_u} + \frac{d}{c_m(g_x)}]$$

Importantly, by dividing Equation S4 by Equation S3, we can then see that the relative frequency of manipulated and unmanipulated reproductives of a given genotype will not change between seasons, and will be equal to:

***Equation S6***

$$\frac{u'_x}{m'_x} = \frac{u_x}{m_x} = \frac{1}{d}[(1-d)\Lambda(g_x) + \frac{c_m(g_x)}{c_u}]$$

If we assume that mutations occur only very rarely we can then say that the resident population will be at its stable dynamics when a mutant invades. We can find the resident's stable dynamics when $N'_2 = 0$, as when $N'_1 = N_1 = m_1 + u_1$. Since $\frac{u_1}{m_1}$ will remain constant, we can then also say that $m'_1 = m_1$, and $u'_1 = u_1$. Via substituting rearrangements of Equation S6 into Equations S3 and S4 we can then obtain:

***Equation S7***

$$m_1 = \frac{nd[c_m(g_1)\gamma_u + c_u\gamma_u(1-\mathrm{d})\Lambda(g_1) + dc_u\gamma_m(g_1)z(g_1)]}{c_m(g_1) + c_u[(1-d)\Lambda(g_1) + dz(g_1)]}$$

***Equation S8***

$$u_1 = \frac{n[c_m(g_1) + c_u(1-d)\Lambda(g_1)][c_m(g_1)\gamma_u + c_u\gamma_u(1-\mathrm{d})\Lambda(g_1) + dc_u\gamma_m(g_1)z(g_1)]}{c_m(g_1) + c_u[(1-d)\Lambda(g_1) + dz(g_1)]}$$

If our mutant genotype is very rare in the population (as we would expect at the point of invasion in an infinitely large population), we can then find $N_2'$ under invasion conditions by substituting Equations S7 and S8 into Equation S5, and taking the 1$^{st}$ order approximation of a taylor series around the point $N_2 = 0 = m_2 + u_2$ (with $m_2 = 0$ and $u_2 = 0$). The 0$^{th}$ order approximation of $N_2'$ will resolve as 0, as such we can write the 1$^{st}$ order approximation as:

$$N_2' = u_2 \frac{\partial N_2'|_{u_2=0\ \&\ m_2=0}}{\partial u_2} + m_2 \frac{\partial N_2'|_{u_2=0\ \&\ m_2=0}}{\partial m_2}$$

We can then take our 1$^{st}$ order approximation and divide it by $N_2$ to find our invasion fitness. Utilising Equation S6 again, we can then find:

***Equation S9***

$$\frac{N_2'}{N_2} = \frac{c_m(g_1)[C_m(g_2)\gamma_u + c_u\gamma_u(1-d)\Lambda(g_2) + c_u d\gamma_m(g_2)z(g_2)]}{c_m(g_2)[c_m(g_1)\gamma_u + c_u\gamma_u(1-d)\Lambda(g_1)\ +\ c_u d\gamma_m(g_1)z(g_1)]}$$

We expect a mutant for manipulation to invade when $\frac{N_2'}{N_2} > 1$ (since this is fitness in discrete time). As such, from Equation S9 we can gain the invasion criteria of:

***Equation S10***

$$\frac{\gamma_u(1-d)\Lambda(g_2) + d\gamma_m(g_2)z(g_2)}{c_m(g_2)} > \frac{\gamma_u(1-d)\Lambda(g_1)\ +\ d\gamma_m(g_1)z(g_1)}{c_m(g_1)}$$

From Equation S10 we can then regain Equations 1 and 2 from the main text, where:

$$\boldsymbol{v_w'} = \gamma_u\Lambda(g_2)$$

$$\boldsymbol{v}_{\boldsymbol{r}}' = \gamma_m(g_2)z(g_2)$$

$$\boldsymbol{c}' = c_m(g_2)$$

$$\boldsymbol{v}_{\boldsymbol{w}} = \gamma_u \Lambda(g_1)$$

$$\boldsymbol{v}_{\boldsymbol{r}} = \gamma_m(g_1)z(g_1)$$

$$\boldsymbol{c} = c_m(g_1)$$

*Model 2 (plastic dispersal)*

We can then derive Equation 4 via the addition of parameter $\alpha(g_x)$. Here $\alpha(g_x)$ is very similar, but not directly equivalent, to our previous parameter of $\boldsymbol{a}$. It fulfils the same function as $\boldsymbol{a}$ while also being a function of foundress genotype, and hence allowing for offspring to respond differently to differing extents of manipulation. When brood 1 offspring change their dispersal behaviour in response to changes in the manipulation to their morphology we can instead write:

***Equation S11***

$$m_x' = \frac{n[u_x\gamma_u + m_x z(g_x)\gamma_m(g_x)]}{u_1 + u_2 + z(g_1)m_1 + z(g_2)m_2} \left[\frac{d(1-\alpha(g_x))}{c_m(g_x)}\right]$$

***Equation S12***

$$u_x' = \frac{n[u_x\gamma_u + m_x z(g_x)\gamma_m(g_x)]}{u_1 + u_2 + z(g_1)m_1 + z(g_2)m_2} \left[\frac{1}{c_m(g_x)}(1-d[1-\alpha(g_x)])\Lambda(g_x) + \frac{1}{c_u}\right]$$

***Equation S13***

$$N_x' = \frac{n[u_x\gamma_u + m_x z(g_x)\gamma_m(g_x)]}{u_1 + u_2 + z(g_1)m_1 + z(g_2)m_2} \left[\frac{1}{c_m(g_x)}(1-d[1-\alpha(g_x)])\Lambda(g_x) + \frac{1}{c_u} + \frac{d(1-\alpha(g_x))}{c_m(g_x)}\right]$$

Following the same steps as4 the derivation of model 1, we can then use Equations S11-S13 to subsequently gain:

$$\frac{u'_x}{m'_x} = \frac{u_x}{m_x} = \frac{1}{d(1-\alpha(g_x))}[(1-d[1-\alpha(g_x)])\Lambda(g_x) + \frac{c_m(g_x)}{c_u}]$$

$$m_1 = \frac{nd[c_m(g_1)\gamma_u + c_u\gamma_u(1-\mathrm{d}[1-\alpha(\mathrm{g}_1)])\Lambda(g_1) + d(1-\alpha(\mathrm{g}_1))c_u\gamma_m(g_1)z(g_1)]}{c_m(g_1) + c_u[(1-d[1-\alpha(\mathrm{g}_1)])\Lambda(g_1) + d(1-\alpha(\mathrm{g}_1))z(g_1)]}[\frac{d(1-\alpha(g_1))}{c_m(g_1)}]$$

$$u_1 = \frac{n[c_m(g_1)\gamma_u + c_u\gamma_u(1-\mathrm{d}[1-\alpha(g_1)])\Lambda(g_1) + d(1-\alpha(g_1))c_u\gamma_m(g_1)z(g_1)]}{c_m(g_1) + c_u[(1-d[1-\alpha(g_1)])\Lambda(g_1) + d(1-\alpha(g_1))z(g_1)]}[\frac{1-d(1-\alpha(g_1))}{c_m(g_1)}\Lambda(g_1) + \frac{1}{c_u}]$$

Enabling us to then gain invasion fitness:

$$\frac{N'_2}{N_2} = \frac{c_m(g_1)[C_m(g_2)\gamma_u + c_u\gamma_u(1-d[1-\alpha(g_2)])\Lambda(g_2) + c_u d[1-\alpha(g_2)]\gamma_m(g_2)z(g_2)]}{c_m(g_2)[c_m(g_1)\gamma_u + c_u\gamma_u(1-d[1-\alpha(g_1)])\Lambda(g_1) + c_u d[a-\alpha(g_1)]\gamma_m(g_1)z(g_1)]}$$

And finally our conditions for invasion:

***Equation S14***

$$\frac{\gamma_u(1-d[1-\alpha(g_2)])\Lambda(g_2) + d[1-\alpha(g_2)]\gamma_m(g_2)z(g_2)}{c_m(g_2)} > \frac{\gamma_u(1-d[1-\alpha(g_1)])\Lambda(g_1) + d[1-\alpha(g_1)]\gamma_m(g_1)z(g_1)}{c_m(g_1)}$$

From which we can regain Equation 4 from the main text, with the slight variation of using $\alpha(g_x)$ instead of $\boldsymbol{a}$.

From the above derivations, an important point is also made clear. If manipulation cannot be limited to those offspring which are less likely to disperse as reproductives (which could be achieved by offspring responding to manipulation), or if all offspring are equally likely to disperse as reproductives, only worker morphological traits which sufficiently increase the productivity of workers relative to the decrease they cause in the productivity of reproductives (such that they're favoured to be applied to all offspring, rather than none) can invade. Consequently, morphological castes could never be expected to evolve this way -since if all offspring are manipulated, the population will end up just as morphologically uniform as when manipulation does not occur.

It is also worth noting that the main limitation of our derivation here is that it does not consider the effects of male production. If the sex allocation strategy in the population diverges from the optimum, the results of our haploid model will then diverge from those of a diploid or haplodiploid model (see main text section 2.2). We can regain the generality of our haploid model by making the additional assumption that mutations for sex allocation are much more frequent than mutations for offspring manipulation.
